# Rapid Cell-Free Combinatorial Mutagenesis Workflow Using Small Oligos Suitable for High-Iteration, Active Learning-Guided Protein Engineering

**DOI:** 10.1101/2025.06.18.660386

**Authors:** Ryan Godin, Sepehr Hejazi, Bret Lange, Basmala Aldamak, Nigel F. Reuel

## Abstract

Active learning-guided protein engineering e2iciently navigates the challenging fitness landscape by screening designs iteratively in a model-guided design-build-test-learn cycle. However, while high iterations boost performance, current workflows reliance on tedious and costly cell-based cloning and expression steps limits the iterations they can practically implement. To address this problem, we present a novel combinatorial mutagenesis workflow that uses small (∼20-40 bp) mutagenic annealed-oligo fragments and cell-free expression to rapidly and conveniently screen protein variants in <9 hours. Using bulk-prepared mutagenic oligos eliminates the need for cloning, PCR-based mutagenesis, or ordering costly genes each screening round. Their >80% size reduction from current fragment-based shu2ling strategies also helps avoid including multiple mutations on the same fragment, reducing the number one must order to cover the design space. By screening 3-10 fragment assemblies for two di2erent proteins, we show our approach is a general, scalable, and cost-e2ective platform for high-iteration protein engineering.

## Introduction

Protein engineering has led to the development of improved therapies^1–3^, sustainable catalysts^4–6^, novel biomaterials^7–9^, and many other important biotechnology applications that impact society. Unfortunately, the process of finding novel protein sequences with enhanced functions remains challenging and prone to failure for several reasons. First, the design space is extremely vast and most sequences in the fitness landscape have little to no function. Just a 10 amino acid protein has 20^10^ possible designs which cannot be practically screened with existing technologies. Even ultrahigh-throughput screening methods can currently screen a small fraction of the possible protein design space at 10^6^-10^8^ sequences per day^10, 11^. Compounding the difficult navigation through the vast design space is the fact that mutations at distant sequence sites may interact in unexpected ways, positive or negative. These epistatic interactions make rational protein engineering difficult to implement and cause the protein fitness landscape to be extremely rugged with small high activity “peaks” surrounded by large regions of inactivity.^12–15^ These regions of inactivity serve as barriers to navigating the fitness landscape and can cause traditional “greedy optimization” strategies like directed evolution to become trapped at a local optimum.^16–18^

To address these issues, researchers have begun leveraging supervised machine learning algorithms in recent years to help navigate the challenging protein design space^16, 17, 19–21^. These models are trained on sequence-function data so they can learn the underlying fitness landscape and identify high-performing regions of the design space for researchers to screen. Machine learning models have been used to engineer a wide variety of protein properties over the years including enzyme activity^8, 16, 20, 22^, thermostability^23^, and binding affinity^24–26^. However, the performance of these models is limited by the quality of the training data on which they are trained,^14, 19, 27^ and it can be difficult to generate a sufficiently informative dataset for model training^14^. This is especially true when only low to medium throughput (10^1^-10^2^) assays exist for the property that is being engineered.^22, 28^ For this reason, active-learning approaches^16, 17, 20, 21, 23^ to protein engineering have become increasingly common since they increase the information density of the training set relative to the number of variants screened^22^. The training data is acquired following a classic design-build-test-learn cycle where protein variants are constructed, their fitness is assayed, the model is trained on this fitness data, and then the model selects the sequences expected to be most informative for learning to be made in the next round. In this way, active learning iteratively constructs a highly informative training set and, therefore, it performs best when subjected to more frequent learning cycles relative to the number of sequences tested^17, 22, 23, 29^.

Unfortunately, recently published workflows using active learning guided design employ lower throughput methods to generate the protein variants for screening. This makes performing each round of variant screening costly and discourages performing more than a few rounds of learning (up to 2-5 rounds in current works)^16, 17, 21, 22^. The slower nature of these mutant generation methods stems from their reliance on cell-based cloning^16, 20–22^ methods which are tedious and add significant time to the process of generating new protein variants between rounds. Some workflows further rely on cell-based expression^16, 17, 22^ which further adds time to the screening process compared to rapid cell-free protein synthesis^30^. Thus, there is a need for active learning-guided protein engineering workflows with more rapid design-build-test-learn cycles, specifically with regards to the creation of DNA templates encoding the variant proteins. This has been partly addressed by a recent report^23^ that utilizes cell-free DNA assembly and expression to rapidly generate and screen glycoside hydrolase variants in 9 hours as part of an automated robotic system. However, it focuses on the creation of glycoside hydrolase variants through the swapping of large sequence domains (sizes of 300-600 bp) derived from natural sequence segments or computational design. While this is a powerful technique for sampling broad sequence diversity, it does not apply to engineering campaigns that target smaller specific amino acid residues (“hotspots”)^31–33^ especially when they are close in sequence space. It is still common practice to rely on commercial, full gene fragment synthesis^16^ to address these higher density mutants which does not meet the cost and time constraints for high-iteration design. Thus, there is still a need for a rapid protein screening workflow compatible with site-specific mutations for active learning-guided design that omits time-consuming cell-based cloning and expression methods.

To address this need, we present a novel workflow for carrying out site-specific combinatorial mutagenesis and protein screening using small mutagenic annealed-oligos and rapid cell-free expression (Figure 1). Our workflow relies on the combinatorial assembly of small, annealed oligo fragments each containing a mutation for a specific residue. These annealed oligos are cost-effective^34^ and straightforward to prepare for Golden Gate assembly reactions. Their small size also increases the number of mutations that can be simultaneously mutated in a region of DNA without needing to include them on the same fragment. This prevents combinatorial coupling and the resulting exponential increase in unique oligos needed to cover the design space. Combined with rapid cell-free expression^35–37^, our workflow is convenient and allows the screening of protein variants from mutagenic fragments in <9 hours. Thus, it is well-suited to streamline protein screening campaigns and enable rapid, high-iteration active learning-guided protein engineering by reducing the cost of performing a screening round. We demonstrate this workflow by using DNA assemblies of varying complexity for both a fluorescent reporter protein and a therapeutically relevant enzyme.

**Figure 1.**
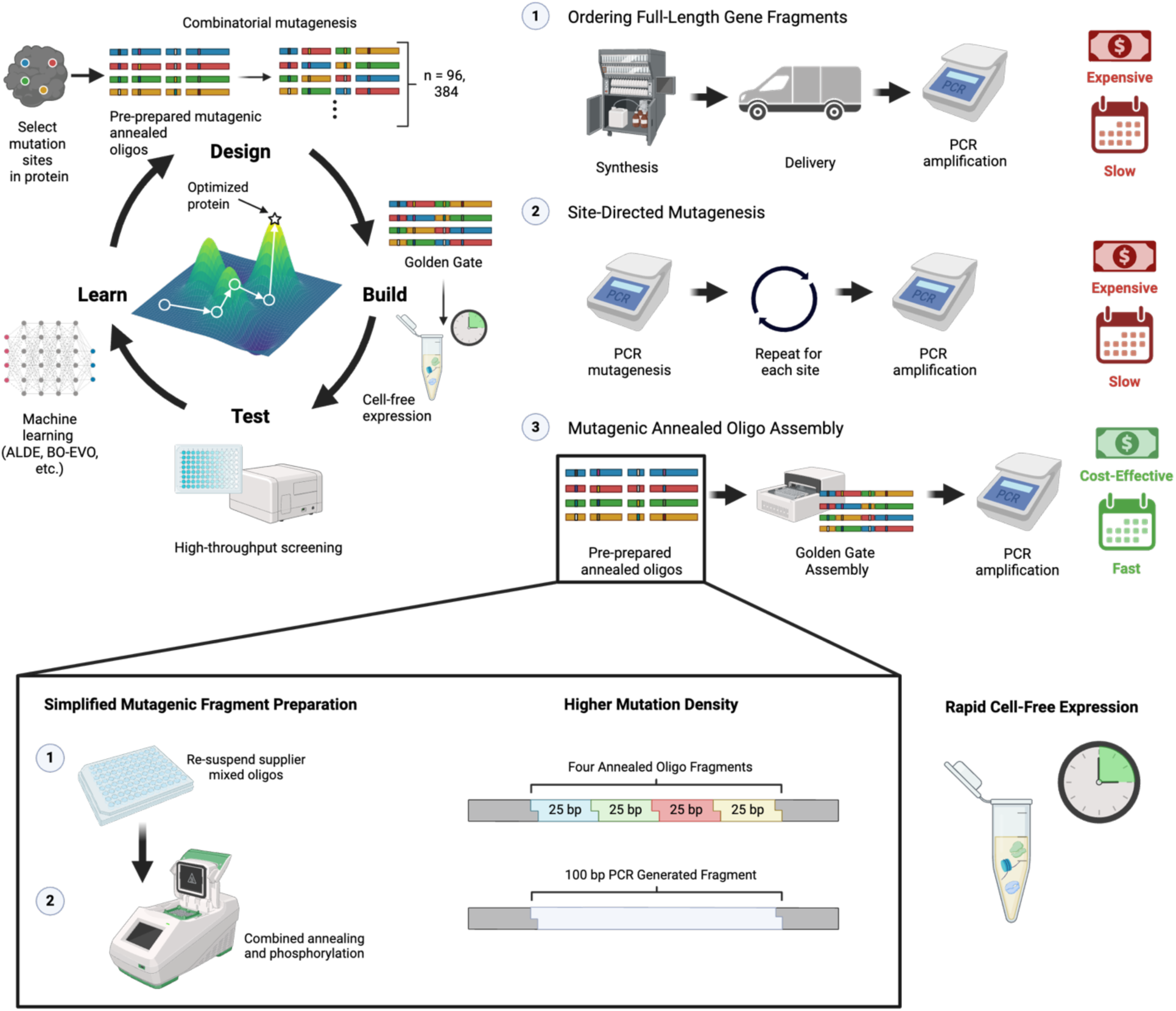
Rapid Combinatorial Mutagenesis Workflow for Active Learning-Guided Design. (A) Overview of proposed cell-free combinatorial mutagenesis workflow and (B) its benefits. The workflow begins by identifying target sites for combinatorial mutagenesis using prior screening data or computational tools (e.g. hotspot predictors). The DNA sequence is then broken up into fragments so that each target residue is contained in a small DNA fragment. Mutant versions of the fragments are then ordered from commercial vendors like IDT and conveniently phosphorylated and annealed into functional mutagenic DNA fragments for golden gate assembly. These DNA sequences are then rapidly expressed into protein variants by cell-free expression, the variants are screened, and their functional data is fed into a machine learning algorithm that determines the next sequences to build and test. This workflow is repeated using the same oligo building blocks, until a protein variant with the desired performance is identified. The benefits of the workflow stem from its use of cell-free expression, absence of cell-based cloning, and use of small annealed oligos as mutagenic DNA fragments. These annealed oligos are simple, cost-effective to prepare in bulk as a library of building blocks using a combined annealing and phosphorylation reaction compared to generating them by site-directed mutagenesis or ordering them as larger gene fragments.

## Results

### Annealed Oligo Assembly Fidelity is Reliable

Since our workflow omits cell-based cloning techniques to enable more rapid design-build-test-learn cycles, inherent quality controls like cell-based selection for replicable plasmids is also absent^38^. Thus, we first evaluated if our annealed oligo assembly method had sufficient fidelity for cell-free screening. This was done by making multiple mutants of a superfolder green fluorescent protein (sfGFP) and a uricase protein and evaluating assembly accuracy with Sanger sequencing and gel electrophoresis. Specifically, we used a three annealed oligo (3-oligo) assembly for sfGFP and 5-oligo assembly for uricase since 3-5 mutation sites is common during combinatorial mutagenesis (Fig 2A, B). We made mutations to Y66 for sfGFP since the resulting color change (blue for Y66H and cyan for Y66W)^39^ is easily assayed with a plate reader while the uricase mutations D44V, Q268R, E279K, K285Q, and H300R were identified from a previous study^40^ to strongly impact specific activity. The agarose gel showed a single band for all mutants analyzed, consistent with the full-length IDT gBlock control except when an oligo was intentionally omitted from the assembly as a negative control (Figures 2C,D). The Sanger sequencing data also confirmed the correct mutations were introduced consistently over multiple replicates demonstrating that our assembly method is reliable (Figures 2E,F and Supporting Information, Figure S1).

**Figure 2.**
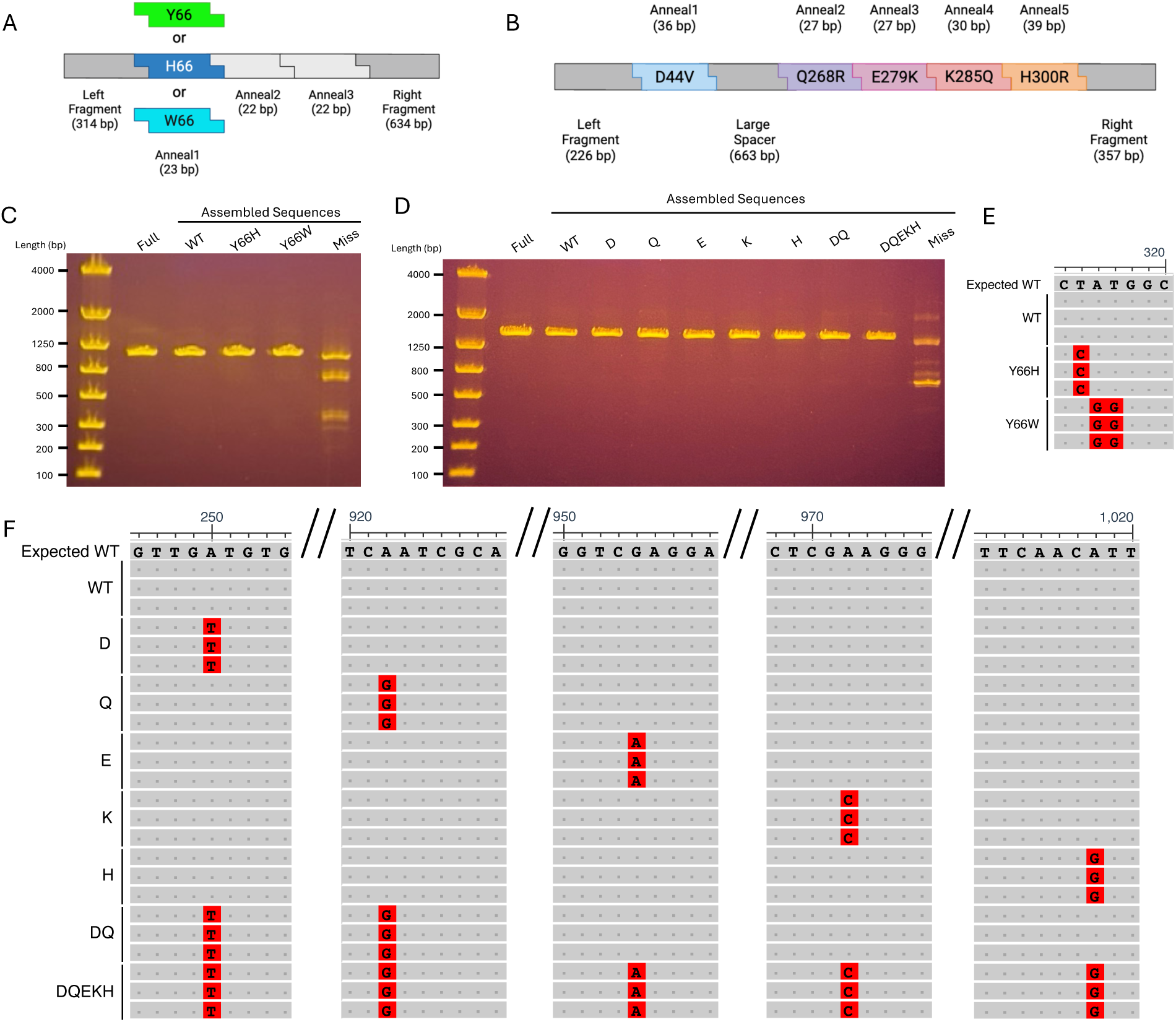
Annealed-Oligo Based Assembly is Reliable. (A,B) Overview of the annealed oligo assemblies used to construct mutants of sfGFP (A) and uricase (B). (C,D) Agarose gel images of the annealed-oligo assemblies for various mutants following PCR-amplification and purification for sfGFP (C) and uricase (D). The “Full” sample is a PCR-amplified IDT gBlock of the full-length non-assembled target gene sequence. The missing oligo assembly “Miss” is the wild-type assembly minus the Anneal1 oligo for sfGFP and Anneal3 oligo for uricase. Gel images are representative of n = 3 independent replicates. (E,F) Multiple sequence alignment of Sanger sequencing data for the sfGFP (E) and uricase (F) mutants showing substitutions at the target mutations sites for multiple mutants and n = 3 independent assemblies.

### Crude PCR-Amplified Assembly Products Express Functional Protein

Having confirmed that we can reliably generate accurate mutant DNA templates with our oligo-based combinatorial mutagenesis strategy, we next tested if functional proteins are expressed from crude PCR-amplified assembly products to save time during screening by omitting DNA purification. We first expressed the sfGFP mutants in a rapid 4-hour cell-free expression reaction and measured their fluorescence at green (Ex/Em: 485/528 nm), blue (Ex/Em: 380/440 nm), and cyan wavelengths (Ex/Em: 440/470 nm) confirming the color change mutations were applied (Figure 3A, B). We also expressed a purified full-length IDT gBlock control and PCR-amplified assembly product at 5 nM and found after controlling for signal saturation (Supporting Information, Figure S2) their yields were not statistically different. For the uricase mutants, we first expressed and purified them to confirm that they were the expected size of ∼ 41 kDa with capillary electrophoresis (Figure 3C). We next assayed the uricase mutants’ ability to degrade uric acid by measuring the decrease in absorbance over time at 293 nm given uric acid’s strong absorption at that wavelength. This activity was normalized by expression level to serve as a proxy for purified specific activity by also measuring the fluorescence produced from a small GFP11-tag with the split GFP assay system^41–43^ (Figure 3D).

**Figure 3.**
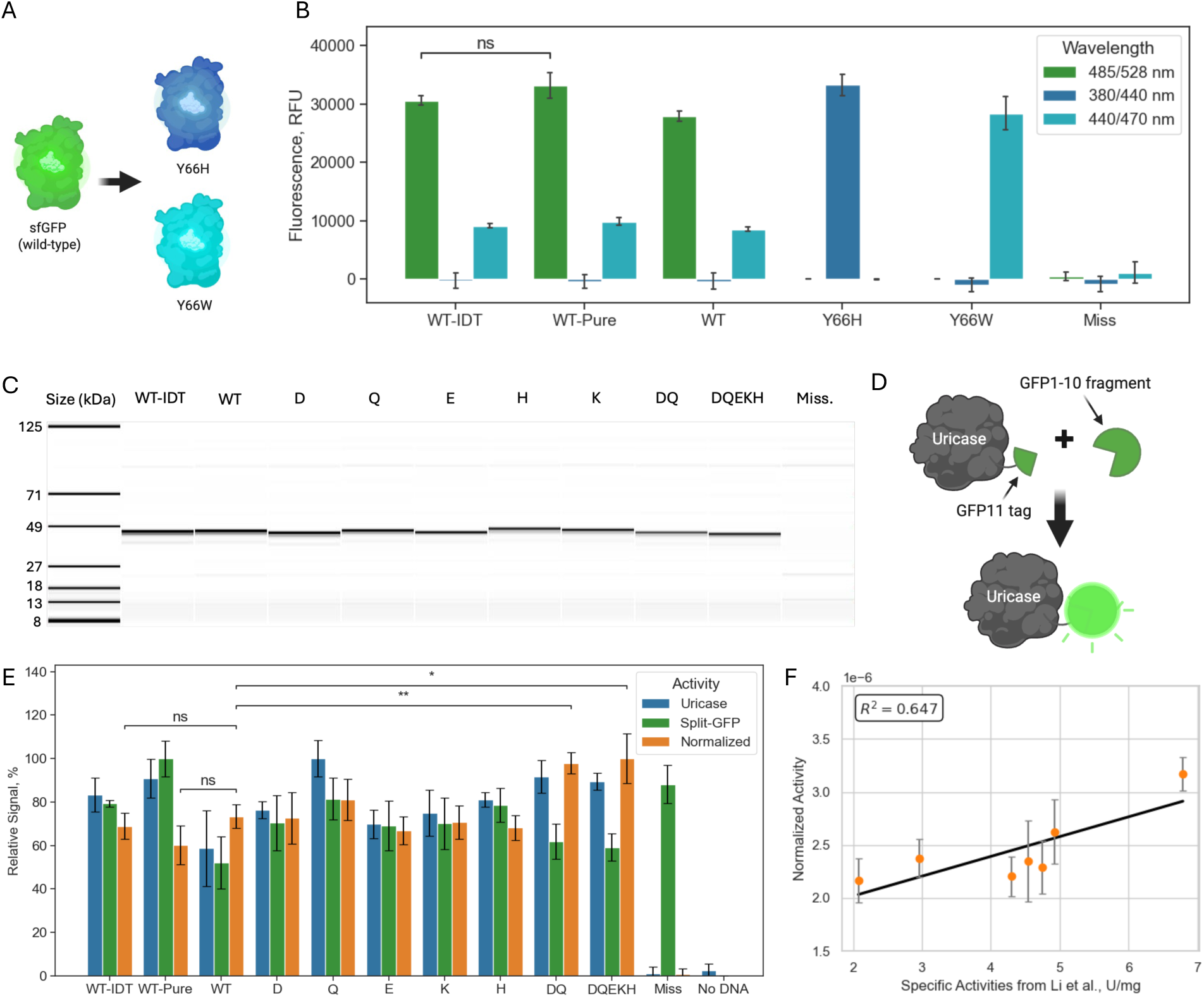
Anneal-Oligo Based Combinatorial Mutagenesis Strategy Produces Functional Protein Variants. (A) Overview of sfGFP mutants and the corresponding color changes that will be assayed for to confirm expression of the variant proteins constructed using the annealed oligo-based assembly strategy. (B) Fluorescence data for the sfGFP protein mutants screened directly in lysate. WT-IDT is the purified full-length gBlock added to the cell-free expression reaction at 5 nM while WT-Pure is the purified assembly also loaded at 5 nM. The rest of the samples were setup using crude PCR product added at 20% (v/v). The wavelengths provided for the fluorescence measurements represent the excitation/emission wavelengths and were taken at different gains so the 485/528, 380/440, 440/470 nm measurements cannot be compared between each other. (C) Capillary electrophoresis data for different purified uricase protein samples expressed from DNA templates constructed using the annealed oligo-based assembly strategy. (D) Diagram of the GFP11-tag used to quantify protein concentration directly in lysate with the split GFP system. This allowed protein activities calculated with lysate-based screens to be normalized to protein amount and minimize biases from expression level differences. The GFP11-tag interacts with a GFP1-10 fragment in the assay’s detector solution to form a functional GFP protein. (E) Scaled activity data for the uricase variants. The uricase activity data represents the ΔAbs(293 nm)/min given uric acid’s strong absorbance at that wavelength. The split GFP data represents the fluorescence stemming from the GFP11-tag and serves as a proxy for expression level. The normalized activity is a proxy for purified specific activity and is the uricase activity divided by the split GFP activity. (F) Regression plot showing the correlation of the normalized activity values measured in this study for the uricase mutants to purified specific activity values (U/mg) reported previously^40^. All mutants except DQEKH were included since the specific activity of this mutant was not reported. All data represents the average of n = 3 independent replicates. Error bars represent the standard deviation from the mean. Statistical significance was calculated using an unpaired student’s t-test (ns = p ≥ 0.05; * = p < 0.05, * * = p < 0.01).

We found that normalized activities of wild-type protein expressed from a full-length IDT gBlock, purified assembly product, or unpurified assembly product were not significantly different further supporting our methods reliability (Figure 3E). While our data also corroborates a previous finding that the DQ mutant has a higher specific activity than the wild-type^40^, the other mutants D, Q, E, K, H were not found to be significantly different despite an overall moderate to strong correlation with the prior report (Figure 3F). Interestingly, we also observed high split GFP signal in the missing oligo assembly despite the absence of uric acid degradation or the presence of a clear truncation product in our capillary electrophoresis data. This may indicate that there is a truncation product containing the split GFP-tag that is smaller than the molecular weight cut-off (MWCO) of the desalting column (7K MWCO) or concentrator (10K MWCO) used during purification. Regardless, our color change data, discrimination of uricase mutants, and general agreement with previous uricase screening data clearly demonstrate that we can use unpurified PCR-amplified oligo assemblies in cell-free reactions to rapidly and reliably screen protein variants.

### Proposed Workflow is Compatible with Automation and High-Throughput Screening

To ensure our cell-free combinatorial mutagenesis strategy was compatible with laboratory automation and high-throughput screening, we simulated a round of screening for the 5-oligo uricase assembly. Specifically, we assayed a common batch size of 96 samples consisting of 6 replicates of 14 mutants, a missing oligo assembly, and a no DNA control. Our simplified screening workflow (Figure 4A) could be completed in < 9 hours using the low-cost Opentrons OT-2 liquid handler for pipetting and reagent transfer resulting in significant time savings during protein engineering. From our screening data, several of the uricase mutants showed significantly different activities from the wild-type (Figure 4B). Replicates were also relatively consistent with variation likely stemming from well-established cell-free variability^44, 45^, difficulty pipetting small μL volumes with the OT-2, and uricase assay noise. While normalization with the GFP11-tag may help minimize some of this well-to-well variability,^42^ signal normalization with the split GFP system was not performed since its long 18–24-hour incubation would significantly slow screening. We did find that using the fluorogenic Janelia Fluor® 646 HaloTag® ligand with HaloTag® system could reduce normalization time to 20-40 min (Supporting Information, Figure S4), but the tags large size may interfere with enzyme activity^42^.

**Figure 4.**
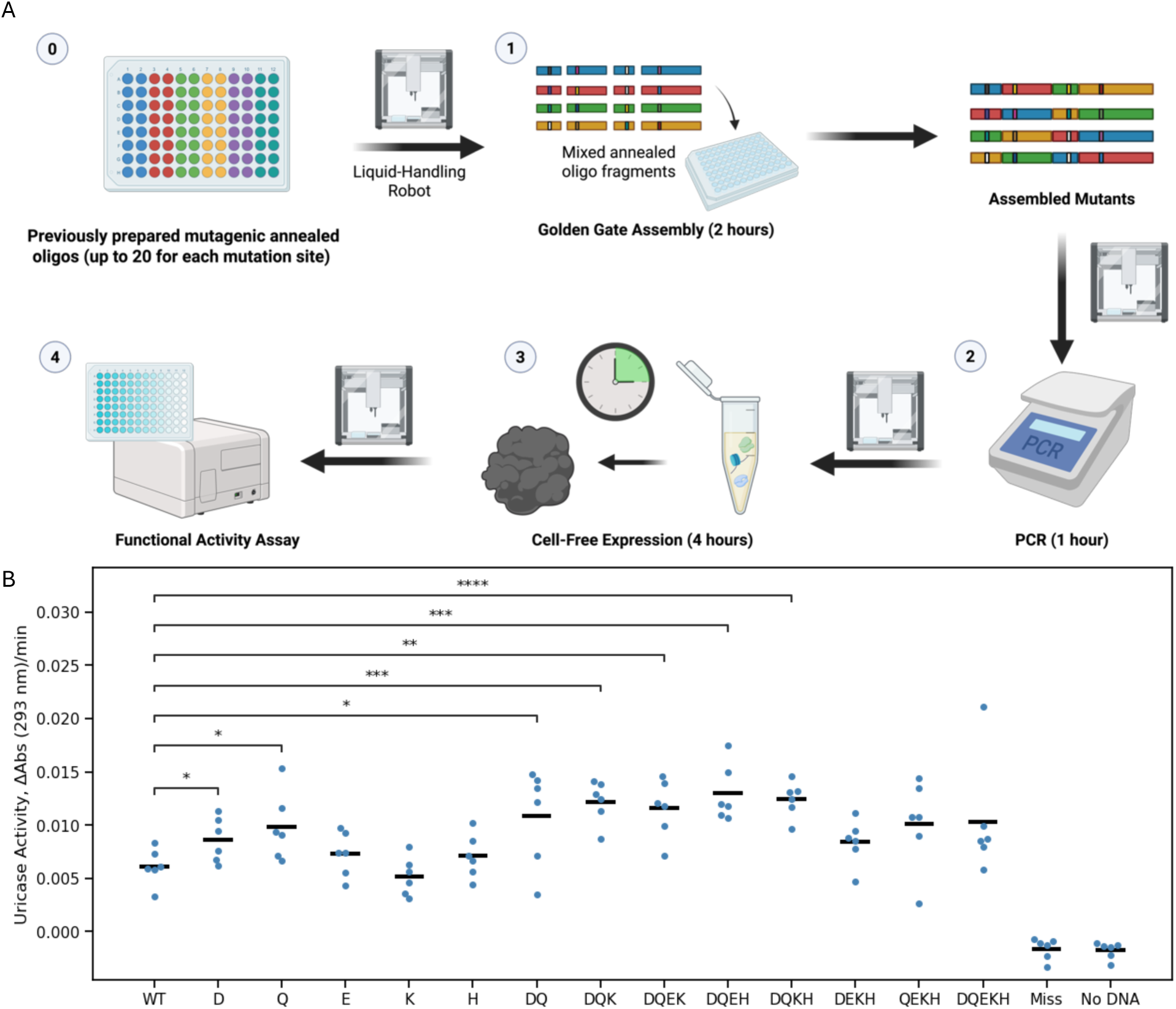
Annealed Oligo Workflow is Compatible with High-Throughput Screening. (A) Overview of simplified screening workflow using cell-free expression and annealed oligos. The workflow can be completed in less than 9 hours which enables very rapid learning cycles for machine-learning guided design. This stems from the simple bulk preparation of annealed oligos prior to screening, cell-free cloning, and rapid cell-free expression. To test this workflow, 96 samples (6 replicates of 16 variants) were assembled and their (B) activities were screened to determine if they were consistent. The black bars represent the average of the six replicate measurements. Statistical significance was calculated using an unpaired student’s t-test (ns = p ≥ 0.05; * = p < 0.05, ** = p < 0.01, *** = p < 0.001, **** = p < 0.0001).

### Annealed Oligo Screening Strategy Scales to 10 Mutagenic Fragments

Having established our annealed oligo screening strategies compatibility with laboratory automation and ability to rapidly screen protein variants, we next tested its fidelity in more complex assemblies since the ease of fragment preparation may make higher-order combinatorial mutagenesis at many sites more practical. Thus, we designed a 10-oligo sfGFP assembly (Figure 5A) and assembled and screened it with cell-free. Agarose gel analysis of crude PCR-amplified assembly products shows that the previously used 10 nM assembly conditions were insufficient for proper construction (Figure 5B). Raising the concentration to 40 nM helped, but further improvements were observed by instead adding the annealed oligos in excess at 200 nM. However, presumable misligation products remained as bands with incorrect length. This is supported by Nanopore sequencing, used before to assess cell-free template quality^46^, which shows a significant increase in deletion products in the region containing the annealed oligos (Figure 5C). Regardless, the 200 nM assemblies were confirmed to have the dominant, correct point mutations upon Sanger sequencing (Figure 5D) and showed the corresponding changes in fluorescence (Figure 5E). This implies that even with misligation products, the assembly fidelity for the complex 10-oligo assembly is sufficient for cell-free protein screening. mutations sites and n = 3 independent assemblies. (E) Fluorescence data for the different assemblies. WT-IDT is the purified full-length gBlock added to the cell-free expression reaction at 5 nM while WT-Pure is the purified assembly also loaded at 5 nM. The rest of the samples were setup using crude PCR product added at 20% (v/v). All data represents the average of n = 3 independent replicates. Error bars represent the standard deviation from the mean. Statistical significance was calculated using an unpaired student’s t-test (ns = p ζ 0.05; * = p < 0.05, ** = p < 0.01, *** = p < 0.001).

**Figure 5.**
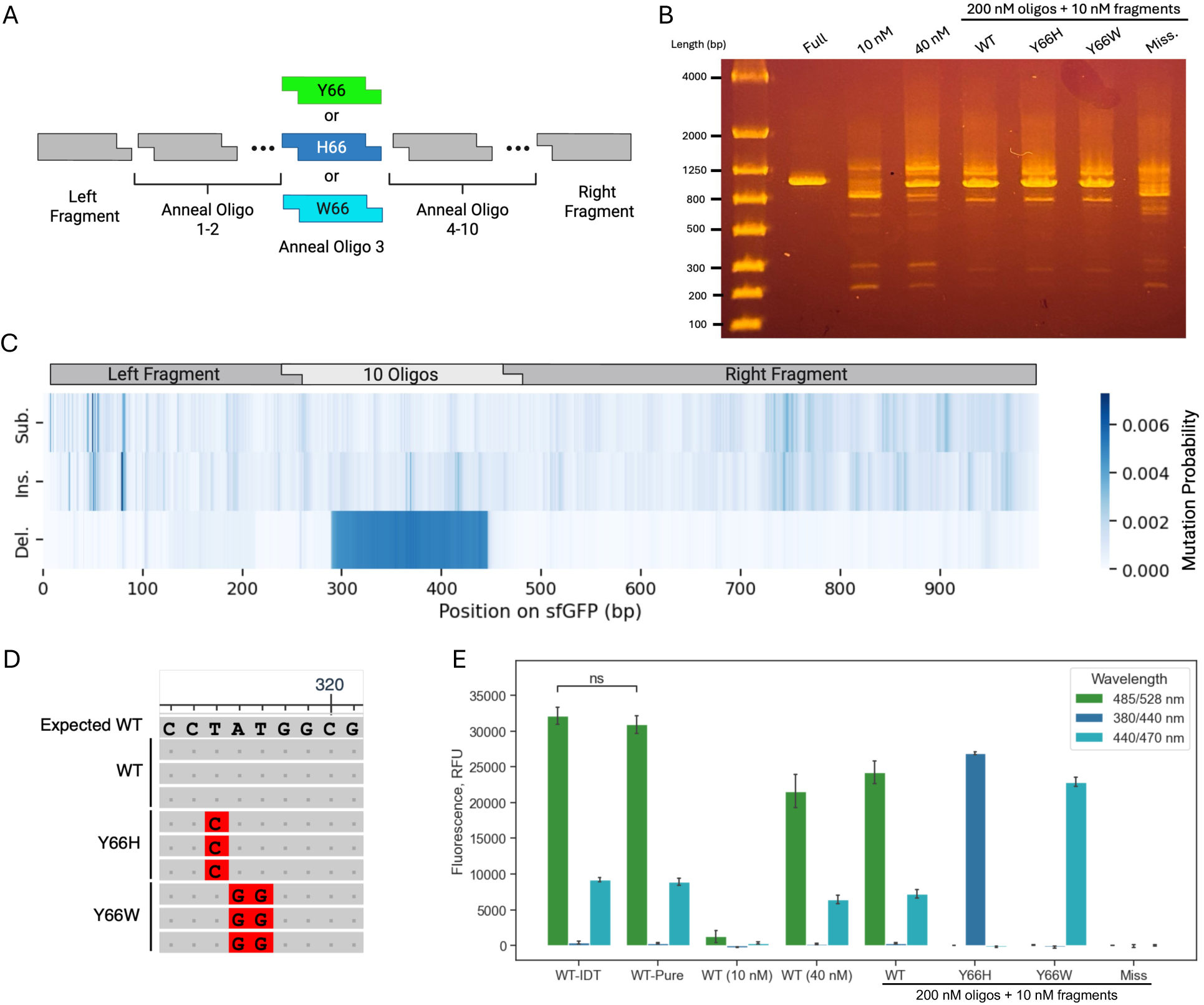
Annealed Oligo-Based Assembly Allows for Complex Assemblies. (A) Overview of 10-oligo sfGFP assembly used to test the scalability of the annealed oligo workflow. (B) Agarose gel showing the difference in assembly fidelity when all fragments/oligos are added to the mix at 10 nM, 40 nM, and when the annealed oligos are added in excess at 200 nM while the fragments are loaded at 10 nM. All samples besides the full are unpurified PCR products. (C) Nanopore sequencing data for the purified 10-oligo assembly showing the probability of substitutions, insertions, and deletions at different positions in the sfGFP LET. (D) Multiple sequence alignment of Sanger sequencing data for the sfGFP mutants showing substitutions at the target mutations sites and n = 3 independent assemblies. (E) Fluorescence data for the different assemblies. WT-IDT is the purified full-length gBlock added to the cell-free expression reaction at 5 nM while WT-Pure is the purified assembly also loaded at 5 nM. The rest of the samples were setup using crude PCR product added at 20% (v/v). All data represents the average of n = 3 independent replicates. Error bars represent the standard deviation from the mean. Statistical significance was calculated using an unpaired student’s t-test (ns = p ^≥^ 0.05; * = p < 0.05, ** = p < 0.01, *** = p < 0.001).

## Discussion

Through sequencing, electrophoretic, and cell-free screening data, we validated a new annealed oligo-based combinatorial mutagenesis workflow for two different proteins and assemblies of varying complexity. Our results show that this workflow is reliable, accurately introducing desired point mutations through the swapping of mutagenic annealed oligo fragments. It is also simple, fast, and convenient, only requiring Golden Gate assembly of the mutagenic annealed oligo fragments, PCR amplification, and rapid cell-free expression before screening. This whole process can be completed in <9 hours and is compatible with laboratory automation. Thus, it will be of interest to both researchers and biofoundries wanting to accelerate their iterative protein engineering campaigns. The ability to conduct a full round of engineering in a single workday makes this workflow especially advantageous to researchers leveraging active learning in their protein engineering campaigns since machine learning models performance appears to benefit from more frequent learning cycles^17, 22, 23, 29^.

Compared to recent mutagenesis workflows utilized in active learning, our annealed oligo-based approach has several distinguishing features. First, it does not rely on any cell-based cloning steps during the DNA assembly process. This significantly accelerates the time required to make the mutant DNA sequences for screening since cell-based cloning adds at least a day to the process^16, 17, 20, 21^ compared to the rapid 3-hour preparation we achieved with Golden Gate and PCR. The second advantage of our workflow is that it uses cell-free expression to rapidly generate protein variants from unpurified PCR-amplified assemblies in 4 hours, adding to the time savings over workflows using cell-based methods. Interestingly, while a few workflows utilizing cell-free expression have been published recently^20, 21^, their use of cell-based cloning techniques significantly extends the time required to generate mutant variants compared to this strategy.

The last major advantage of our workflow for combinatorial mutagenesis stems from the use of annealed oligos as the mutagenic fragments in Golden Gate assemblies. First, the small size of these fragments (15-30 bp) means more can be fit into a region of DNA compared to larger PCR-generated mutagenic DNA fragments. This increases the mutation sites that can be fit into a region of DNA and interchanged independently, minimizing the likelihood of introducing combinatorial complexity by having multiple mutations on the same DNA fragment. As an example, having just two mutation sites on the same DNA fragment requires 400 (20^2^) fragments for complete mutation coverage compared to 40 (2 x 20) if the sites are on individual fragments, a significant cost increase especially as the number of desired mutation sites increase. Secondly, their preparation is simple. The two oligos forming the annealed fragment can be ordered pre-mixed from IDT or mixed manually if ordered from different DNA vendors. Once mixed, the oligos need to be re-suspended if not already, then added to a combined 2-3 hour annealing and phosphorylation reaction. After this, they are ready for direct use in Golden Gate assembly and can be stored in bulk and used over multiple learning cycles. This preparation of a library of DNA assembly blocks is much more practical and cost-efficient than producing each mutagenic fragment using site-directed mutagenesis as previously done^17^, a benefit that scales with the number of mutation sites selected for engineering. Since current machine learning models perform quite well on 4-5 mutation site fitness landscapes, increasing the number of mutation sites may be a worthwhile pursuit to help differentiate model performance and motivate further model advancements. For example, the computation time for models that predict fitness for the whole landscape between each learning round, like Bayesian optimization, scales exponentially with the design space,^17^ potentially becoming limiting in the future as more complex landscapes are explored. Having demonstrated the reliable assembly and screening of a 10-oligo sfGFP mutant, the ease of mutagenic fragment preparation should make our annealed oligo workflow well-suited for studies exploring active-learning model performance on scaled-up design spaces consisting of >7 simultaneous mutation sites.

While our annealed oligo design strategy was shown to be suitable for high-throughput protein screening, there are few limitations that merit further study and innovation. First, although this new workflow should be compatible with active-learning, we acknowledge that machine learning was not yet implemented in this method development study – something that will quickly follow in a subsequent study. Regardless, the workflow should still prove useful in the absence of machine learning to streamline iterative site-specific mutagenesis workflows due to its ability to rapidly generate and screen protein variants with controllable mutations at user-defined sites. Second, even though we demonstrated site-specific mutagenesis with a 10-oligo sfGFP assembly, there were clear misligation products present that could not be eliminated through simple PCR optimization. Although the presence of the misligation products did not appear to affect the screening data based on similar yields for the full-length IDT gBlock and purified assembly, this was not rigorously investigated and cannot be ruled out. Future work attempting to minimize the presence of misligations could explore data-driven design for selecting the Golden Gate overlaps^34, 47, 48^; further tuning the fragment concentrations; or exploring different Golden Gate conditions like cycle number, digestion time, ligation time, and ligation temperature^49^. Additionally, despite not being a limitation of our annealed-oligo workflow specifically, the split GFP system could not be used generally to identify mis-assemblies since it showed high activity when an oligo was absent from the assembly. Thus, it would be ambiguous during screening if a protein variant showed no activity because the introduced mutations render it inactive or if a pipetting error led to an assembly failure. The long incubation time also poses a roadblock to rapid, high-iteration screening. Faster methods for normalization exist,^50–52^ but they suffer from other limitations like non-specific binding highlighting the need for better normalization methods overall. Finally, despite our proposed workflow advantages, it is not a panacea. If the mutation sites are side-by-side, the spacing will be too small for a Golden Gate site and the mutations will need to be included in a single oligo leading to combinatorial coupling and many mutant oligos will therefore be needed to cover the design space. In this situation, other mutational strategies like PCR-based mutagenesis with a single primer may be favored^16^. Additionally, this workflow is only medium throughput (10-100s of variants) due to its reliance on plate-based assembly and expression methods. If the protein of interest is compatible with high- or ultrahigh-throughput screening, then those methods should be strongly considered.

## Conclusion

Despite these limitations, our oligo-based cell-free screening workflow is well-suited to a wide-range of protein engineering tasks. Advantages of the workflow include: (1) speed stemming from its use of cell-free cloning and expression, (2) compatibility with laboratory automation, (3) convenient bulk mutagenic fragment preparation with annealed oligos that don’t require preparation each round, and (4) absence of tedious purification steps during DNA template preparation and also during protein screening if the assay is compatible with lysate.

The workflow’s facile, rapid nature will interest those wanting to streamline their iterative site-specific combinatorial mutagenesis workflows and improve active learning performance by increasing the number of learning cycles performed in a given period of time. The practical nature of the oligo-fragment preparation combined with our use of a low-cost liquid handler should also help improve accessibility to this technique^53^ for machine learning-guided protein engineering and facilitate complex combinatorial mutagenesis campaigns with more simultaneous mutation sites (n > 7) than explored previously. This will enable researchers to test the limits of current active learning-guided protein engineering methods and models. Thus, our workflow is not only well-positioned to streamline the design of improved therapies, biocatalysts, biomaterials, and other protein engineering targets, but it should also advance the use of active learning-guided design in protein engineering.

## Methods

### DNA Sequences and Preparation

All DNA sequences, primers, and oligos used in this study are provided in Supporting Information, Tbls. S1-S4. The DNA sequence encoding the sfGFP protein used in this study is a linear expression template (LET) reported in a previous study^35^. The uricase protein’s amino acid sequence was also taken from a prior report^54^ and expressed from a LET by reverse translating the sequence, codon optimizing it using IDT’s codon optimization tool, and substituting the resulting sequence into the coding region for the sfGFP LET. Larger gene fragments used to express the full-length genes or large spacer regions in the annealed oligo assemblies were ordered as IDT gBlocks. Once delivered, the gene fragments were resuspended in nuclease-free water, PCR amplified with the Q5® High-Fidelity 2X Master Mix based on manufacturer recommendations, purified using the DNA Clean & Concentrator-5 kit (Zymo Research), and the concentration was calculated with the Qubit™ 1X dsDNA High Sensitivity kit (#Q3323, ThermoFisher). The thermocycler parameters used for the PCR reaction are an initial denaturation at 98°C for 3 min; 17 cycles of denaturation at 98°C for 10 s, annealing at 60°C for 30 s, and elongation at 72°C for 30 s; and a final elongation step at 72°C for 2 min before being held at 4°C.

The annealed oligos used as mutagenic gene fragments for combinatorial mutagenesis were designed to have the 4-bp Golden Gate overhangs once annealed. Compatibility of the overhangs used in an assembly was verified with NEB’s NEBridge Ligase Fidelity Viewer® to ensure they had a high fidelity (> 98%). The top and bottom strands were ordered from IDT as RxnReady™ Primer Pools, normalized to 10 nmol with standard desalting purification. Once delivered, the annealed oligos were resuspended to 50 μM with nuclease-free duplex buffer (#11-01-03-01, IDT). They were then annealed and phosphorylated simultaneously based on a previous protocol^55^ by mixing 2 μL of the 50 μM oligos, 2 μL of 10X T4 DNA Ligase Buffer (NEB), 1 μL of T4 Polynucleotide Kinase (NEB), and 15 μL of nuclease-free water. Once assembled, the reaction was carried out on a thermocycler programmed to perform a phosphorylation step at 37°C for 1 h, a heat inactivation step at 95°C for 3 min, and an annealing step consisting of a 10 min incubation with a 0.1°C/s cooling ramp at the following gradually lower temperatures: 85, 75, 65, 55, 45, 35, and 25°C. The annealed oligos were then diluted to the correct concentration before use in Golden Gate assembly as discussed below.

### Annealed-Oligo Based Combinatorial Mutagenesis Workflow

To carry out the combinatorial mutagenesis workflow, we first prepared a “backbone mix” of all the large gBlock spacer fragments that were invariant among the mutant assemblies. All the fragments were loaded into this mixture at 10 nM unless otherwise specified. The backbone mix was then mixed with each annealed oligo (also at 10 nM) required for the assembly. The resulting DNA mix was used to setup a Golden Gate assembly by preparing a solution of: 58.33% DNA mix, 26.67% nuclease-free water, 10% 10X T4 DNA ligase buffer (#B0202, NEB), and 5% Golden Gate enzyme mix (BsmBI-v2: #E1602 for sfGFP and BsaI-HF-v2: #E1601 for uricase, both from NEB). The Golden Gate assembly was then run in the thermocycler using the following parameters: 30 cycles of digestion at 37°C (for BsaI-HF-v2) or 42°C (for BsmBI-v2) for 1 min and ligation at 16°C for 1 min. The cycles were followed by heat inactivation at 60°C for 5 min before being held at 4°C.

Once the Golden Gate assembly was complete, a PCR reaction was setup as above using the crude product as template loaded at 20% (v/v). However, the number of cycles varied from 21 cycles for the 3-oligo and 5-oligo assemblies to 23 cycles for the 10-oligo assembly. The crude PCR products were then expressed rapidly using a 4-hour cell-free expression reaction incubated at 37°C. The cell-free reaction was setup as described previously^56^ with a few modifications. Specifically, tRNA was omitted from the energy mix preparation and the crude PCR-amplified DNA template was added at 20% (v/v). The purified DNA templates were loaded at 5 nM as before.

### Evaluation of Annealed Oligo Assembly DNA Template Quality

DNA template quality for the oligo-based assemblies was evaluated using Sanger sequencing and gel electrophoresis. The PCR-amplified Golden Gate assembly product was purified the same way as the gBlock gene fragments for Sanger sequencing. DNA Gel electrophoresis was performed using the FlashGel® System (Lonza), a 1.2% FlashGel®DNA Cassette (#57023, Lonza), and a 100-4000 bp DNA marker (#50473, Lonza). Samples were prepared with 5X FlashGel® Loading Dye (Lonza) according to the manufacturer recommendations. Purified samples were loaded between 20-25 ng while unpurified samples were loaded at 40% (v/v). The electrophoresis was run at 275 V until the desired separation was achieved.

### Protein Screening

Protein mutants were screened immediately upon completion of their cell-free expression. For the sfGFP variants, 5 μL of the cell-free expression product was added to a low volume, black, flat transparent bottom 384-well plate (#3540, Corning). The plate was sealed, put in a Tecan Spark multimode plate reader heated to 30°C, and fluorescence measurements were taken every minute for 10 minutes with a 40 second shaking at 240 rpm and 20 second waiting period in between. The fluorescence was measured at three different wavelength, all with a bandwith of 10 nm. The first was green at an excitation/emission of 485/528 nm and a gain of 62. The second was blue at an excitation/emission of 380/440 nm and a gain of 100.

The third was cyan at an excitation/emission of 440/470 nm and a gain of 100. The final measurement was used for the fluorescence reading after background subtraction with a no DNA control cell-free expression sample.

For the uricase variants, the uricase activity assay was performed by taking 4 μL of cell-free expression product and mixing with 96 μL of assay buffer consisting of 62.5 μM of uric acid in a 45 mM Tris-HCl buffer at pH 8.5 in a clear, flat bottom, 384-Well UV-Star® microplate (#781801, Greiner). The well plate was then immediately added to the Tecan Spark plate reader pre-heated to 28°C and the kinetic assay was started. Specifically, the plate was shaken for 20 seconds at 360 rpm and allowed to rest for 40 seconds. The absorbance at 293 nm was then measured every 20 seconds for 15 min without shaking. The uricase activity was then calculated from this data by fitting the data between 1-10 min to a linear regression curve and taking the slope as the reaction rate in ΔAbs(293 nm)/min. For the split GFP assay, 2 μL of cell-free expression product was mixed with 78 μL of Fold-N-Glow™ Split GFP Detection Reagent (#21004001, Sandia Biotech) in a black, flat transparent bottom 384-well plate (#242764, Thermo Scientific); shaken at 237 rpm for 5 min; then left to incubate for ∼24 hours at 4°C. After incubation, the plate was added to the plate reader pre-heated to 30°C and fluorescence measurements were taken after 10 min of continuous shaking at 240 rpm followed by a 30 s wait period. This process was repeated for 2 hours with fluorescence readings being taken with an excitation/emission of 485/525 nm, bandwidth of 15 nm, and a gain of 75. The final measurement was used for the fluorescence reading after background subtraction with a no DNA control cell-free expression sample and this reading was taken as a proxy for expression. The normalized activity was obtained by dividing the ΔAbs(293 nm)/min by the split GFP fluorescence measurement.

### Electrophoretic Analysis of Protein Samples

Proteins were first purified before electrophoretic analysis. Protein variants were prepared in cell-free expressions following the combinatorial mutagenesis workflow above. However, the cell-free expressions were incubated at 30°C overnight. After incubation, samples were purified using the Strep-II tag (WSHPQFEK) and the Strep-Tactin®XT 4Flow® high capacity Spin Column Kit (#2-5151-000, IBA Lifesciences) according to manufacturer recommendations. Once purified, a buffer exchange was performed using Zeba™ Spin Desalting Columns, 7K MWCO (#89882, ThermoFisher) to transfer the protein to a 50 mM HEPES, 100 mM NaCl buffer at pH 7.5. The protein samples were then concentrated using a 0.5 mL Pierce™ Protein Concentrators PES, 10K MWCO spin column (#88513, ThermoFisher) and their concentrations were calculated using the Qubit Protein Assay (#Q3321, ThermoFisher). Finally, the protein samples were processed using the ProteinEXact™ HR Assay on the LabChip® GXII Touch according to manufacturer recommendations using 2-mercaptoethanol as the reducing agent and a protein load between 180-230 ng/μL.

### High-Throughput Screening Validation

The protein screening workflow was divided into five stages: fragment mixing, Golden Gate assembly, PCR amplification, cell-free expression, and assay screening. The pipetting and general setup for each stage was performed using the Opentrons OT-2 robot with custom scripts provided in the Supporting Information. The fragment mixing was performed by combining 2 μL each of the backbone mix and required annealed oligos to make a specific mutant in the wells of an Applied Biosystems™ MicroAmp™ Optical 384-Well Reaction Plate (#4309849, ThermoFisher). The Golden Gate script then transferred 3.5 μL of the resulting DNA mixes to the wells of a second plate and mixed them with 2.5 μL of a Golden Gate master mix (made using the above recipe but without the DNA). The plate was then sealed with an adhesive film (#4306311, ThermoFisher) and the reaction was run as previously described in an Applied Biosystems™ VeritiPro™ 384-well Thermal Cycler. Once completed, the PCR amplification script then transferred 2 μL of the resulting Golden Gate product to a new 384-well PCR plate and mixed it with 8 μL of a PCR master mix (25% v/v nuclease-free water, 6.25% v/v forward and reverse primer, and 62.5% Q5 2X master mix). The plate was then sealed and the PCR reaction was run as previously described in the 384-well thermal cycler for 21 cycles.

Afterwards, the cell-free script transferred 2 μL of the crude PCR product to a low volume, black, flat transparent bottom 384-well plate; sealed with an Axygen PCR-SP plate film; and left to incubate for 4 hours at 37°C. Once the cell-free reaction was completed, the assay screening script was run. It transferred 4 μL of completed cell-free reaction and set up the uricase activity assay as above. The plate was then transferred to the plate reader and the ΔAbs(293 nm)/min was measured as previously described.

### Nanopore Sequencing and Analysis

The 10-oligo sfGFP assembly was prepared as mentioned above. However, 3 min digestion and ligation cycles were used for the Golden Gate assembly and the concentration of DNA fragments in the backbone mix was 2.5 nM. This assembly was repeated with different annealed oligo fragment concentrations of 200, 100, 50, and 25 nM for both a 16C and 25C ligation temperature. These samples were then purified after PCR amplification as above but were eluted after purification with IDTE buffer at pH 7.5 (#11-01-02-02, IDT). The samples were sent to Iowa State’s DNA facility for preparation as a pooled, barcoded library and sequencing on an Oxford Nanopore GridION X5. Following a previous report^46^, the raw sequencing files (POD5) were basecalled using Guppy basecaller and aligned to the expected sfGFP reference sequence with MiniMAP2. The CIGAR string of the alignment output was analyzed to extract the mutation rates and positions on the sfGFP reference gene for plotting.

## Supporting Information

Supporting information included with this work are:

- Sequences for all DNA parts used, experimental information about the HaloTag assay setup and associated signal normalization, uncropped multiple sequence alignments for the Sanger sequencing data, calibration data for sfGFP and split GFP signal measurements, and calibration curves for uricase mutants tagged with HaloTag (PDF).
- Scripts needed to perform the automated screening steps with the Opentrons OT-2 robot and custom labware definitions (PY, JSON).

## Acknowledgements

Figures were created with the assistance of BioRender.com. Research reported in this publication was supported by NIGMS of the National Institutes of Health under award number R35GM138265. The content is solely the responsibility of the authors and does not necessarily represent the official views of the National Institutes of Health. RG was also supported by the National Science Foundation Graduate Research Fellowship under Grant No. 2336877.

## Author Contributions

RG planned the study, carried out experiments, analyzed data, and wrote the manuscript. SH analyzed the Nanopore sequencing data. BL performed electrophoretic analysis of the purified protein samples. BA assisted with experiments for the 10-oligo sfGFP assembly. NFR directed the study and edited the manuscript.

## References

1. Lee, C.-H. et al. An engineered human Fc domain that behaves like a pH-toggle switch for ultra-long circulation persistence. Nat Commun 10, 5031 (2019).

2. Ebrahimi, S. B. & Samanta, D. Engineering protein-based therapeutics through structural and chemical design. Nat Commun 14, 2411 (2023).

3. Chennamsetty, N., Voynov, V., Kayser, V., Helk, B. & Trout, B. L. Design of therapeutic proteins with enhanced stability. Proc. Natl. Acad. Sci. U.S.A. 106, 11937–11942 (2009).

4. Rocha, R. A., Speight, R. E. & Scott, C. Engineering Enzyme Properties for Improved Biocatalytic Processes in Batch and Continuous Flow. Org. Process Res. Dev. 26, 1914– 1924 (2022).

5. Athavale, S. V. et al. Biocatalytic, Intermolecular C−H Bond Functionalization for the Synthesis of Enantioenriched Amides. Angew Chem Int Ed 60, 24864–24869 (2021).

6. Coelho, P. S., Brustad, E. M., Kannan, A. & Arnold, F. H. Olefin Cyclopropanation via Carbene Transfer Catalyzed by Engineered Cytochrome P450 Enzymes. Science 339, 307–310 (2013).

7. J., B., M. M., B. & Chanda, K. Evolutionary approaches in protein engineering towards biomaterial construction. RSC Adv. 9, 34720–34734 (2019).

8. Chu, S., Wang, A. L., Bhattacharya, A. & Montclare, J. K. Protein based biomaterials for therapeutic and diagnostic applications. Prog. Biomed. Eng. 4, 012003 (2022).

9. Straley, K. S. & Heilshorn, S. C. Independent tuning of multiple biomaterial properties using protein engineering. Soft Matter 5, 114–124 (2009).

10. Gantz, M., Neun, S., Medcalf, E. J., Van Vliet, L. D. & Hollfelder, F. Ultrahigh-Throughput Enzyme Engineering and Discovery in *In Vitro* Compartments. Chem. Rev. 123, 5571–5611 (2023).

11. Agresti, J. J. et al. Ultrahigh-throughput screening in drop-based microfluidics for directed evolution. Proc. Natl. Acad. Sci. U.S.A. 107, 4004–4009 (2010).

12. Miton, C. M. & Tokuriki, N. How mutational epistasis impairs predictability in protein evolution and design. Protein Science 25, 1260–1272 (2016).

13. Meger, A. T. et al. Rugged fitness landscapes minimize promiscuity in the evolution of transcriptional repressors. Cell Systems 15, 374–387.e6 (2024).

14. Wittmann, B. J., Yue, Y. & Arnold, F. H. Informed training set design enables edicient machine learning-assisted directed protein evolution. Cell Systems 12, 1026–1045.e7 (2021).

15. Papkou, A., Garcia-Pastor, L., Escudero, J. A. & Wagner, A. A rugged yet easily navigable fitness landscape. Science 382, eadh3860 (2023).

16. Yang, J. et al. Active learning-assisted directed evolution. Nat Commun 16, 714 (2025).

17. Hu, R. et al. Protein engineering via Bayesian optimization-guided evolutionary algorithm and robotic experiments. Briefings in Bioinformatics 24, bbac570 (2023).

18. Weinreich, D. M., Delaney, N. F., DePristo, M. A. & Hartl, D. L. Darwinian Evolution Can Follow Only Very Few Mutational Paths to Fitter Proteins. Science 312, 111–114 (2006).

19. Wu, Z., Kan, S. B. J., Lewis, R. D., Wittmann, B. J. & Arnold, F. H. Machine learning-assisted directed protein evolution with combinatorial libraries. Proceedings of the National Academy of Sciences of the United States of America 116, 8852–8858 (2019).

20. Landwehr, G. M. et al. Accelerated enzyme engineering by machine-learning guided cell-free expression. Nat Commun 16, 865 (2025).

21. Thornton, E. L., Boyle, J. T., Laohakunakorn, N. & Regan, L. Cell-Free Protein Synthesis as a Method to Rapidly Screen Machine Learning-Generated Protease Variants. ACS Synth. Biol. 14, 1710–1718 (2025).

22. Vornholt, T. et al. Enhanced Sequence-Activity Mapping and Evolution of Artificial Metalloenzymes by Active Learning. ACS Cent. Sci. acscentsci.4c00258 (2024) doi:10.1021/acscentsci.4c00258.

23. Rapp, J. T., Bremer, B. J. & Romero, P. A. Self-driving laboratories to autonomously navigate the protein fitness landscape. Nat Chem Eng 1, 97–107 (2024).

24. Li, L. et al. Machine learning optimization of candidate antibody yields highly diverse sub-nanomolar adinity antibody libraries. Nat Commun 14, 3454 (2023).

25. Makowski, E. K., Chen, H.-T. & Tessier, P. M. Simplifying complex antibody engineering using machine learning. Cell Systems 14, 667–675 (2023).

26. Frey, N. C. et al. Lab-in-the-loop therapeutic antibody design with deep learning. Preprint at 10.1101/2025.02.19.639050 (2025).

27. Fox, R. et al. Optimizing the search algorithm for protein engineering by directed evolution. Protein Engineering Design and Selection 16, 589–597 (2003).

28. Biswas, S., Khimulya, G., Alley, E. C., Esvelt, K. M. & Church, G. M. Low-N protein engineering with data-edicient deep learning. Nature Methods 18, 389-+ (2021).

29. Azimi, J., Fern, A. & Fern, X. Z. Batch Bayesian Optimization via Simulation Matching. In Advances in Neural Information Processing Systems 23 (2010).

30. Garenne, D. et al. Cell-free gene expression. Nat Rev Methods Primers 1, 49 (2021).

31. van der Meer, J., Biewenga, L. & Poelarends, G. J. The Generation and Exploitation of Protein Mutability Landscapes for Enzyme Engineering. ChemBioChem 17, 1792–1799 (2016).

32. Bao, Y., Xu, Y. & Huang, X. Focused rational iterative site-specific mutagenesis (FRISM): A powerful method for enzyme engineering. Molecular Catalysis 553, 113755 (2024).

33. Erkanli, M. E., El-Halabi, K., Kang, T. K. & Kim, J. R. Hotspot Wizard-informed engineering of a hyperthermophilic β-glucosidase for enhanced enzyme activity at low temperatures. Biotech & Bioengineering 121, 2079–2090 (2024).

34. Hoch, S. Y. et al. GGASSEMBLER: Precise and economical design and synthesis of combinatorial mutation libraries. Protein Science 33, (2024).

35. Dopp, J., Rothstein, S., Mansell, T. & Reuel, N. Rapid prototyping of proteins: Mail order gene fragments to assayable proteins within 24 hours. BIOTECHNOLOGY AND BIOENGINEERING 116, 667–676 (2019).

36. Dopp, J. & Reuel, N. Simple, functional, inexpensive cell extract for in vitro prototyping of proteins with disulfide bonds. BIOCHEMICAL ENGINEERING JOURNAL 164, (2020).

37. Dopp, B. J. L., Tamiev, D. D. & Reuel, N. F. Cell-free supplement mixtures: Elucidating the history and biochemical utility of additives used to support in vitro protein synthesis in E. coli extract. Biotechnology Advances 37, 246–258 (2019).

38. Chao, R., Yuan, Y. & Zhao, H. Recent advances in DNA assembly technologies. FEMS Yeast Res n/a-n/a (2014) doi:10.1111/1567-1364.12171.

39. Heim, R., Prasher, D. C. & Tsien, R. Y. Wavelength mutations and posttranslational autoxidation of green fluorescent protein. Proc. Natl. Acad. Sci. U.S.A. 91, 12501– 12504 (1994).

40. Li, W. et al. Directed evolution to improve the catalytic ediciency of urate oxidase from Bacillus subtilis. PLoS ONE 12, e0177877 (2017).

41. Cabantous, S., Terwilliger, T. & Waldo, G. Protein tagging and detection with engineered self-assembling fragments of green fluorescent protein. NATURE BIOTECHNOLOGY 23, 102–107 (2005).

42. Santos-Aberturas, J., Dörr, M., Waldo, G. S. & Bornscheuer, U. T. In-Depth High-Throughput Screening of Protein Engineering Libraries by Split-GFP Direct Crude Cell Extract Data Normalization. Chemistry & Biology 22, 1406–1414 (2015).

43. Yuan, Q. et al. Rapid prototyping enzyme homologs to improve titer of nicotinamide mononucleotide using a strategy combining cell-free protein synthesis with split GFP. Biotech & Bioengineering bit.28326 (2023) doi:10.1002/bit.28326.

44. Rhea, K. A. et al. Variability in cell-free expression reactions can impact qualitative genetic circuit characterization. Synthetic Biology 7, ysac011 (2022).

45. Cole, S. D. et al. Quantification of Interlaboratory Cell-Free Protein Synthesis Variability. ACS Synth. Biol. 8, 2080–2091 (2019).

46. Hejazi, S., Ahsan, A., Kashani, S., Tameiv, D. & Reuel, N. F. Amplified DNA heterogeneity assessment with Oxford Nanopore sequencing applied to cell free expression templates. PLoS ONE 19, e0305457 (2024).

47. Pryor, J. M., Potapov, V., Bilotti, K., Pokhrel, N. & Lohman, G. J. S. Rapid 40 kb Genome Construction from 52 Parts through Data-optimized Assembly Design. ACS Synth. Biol. 11, 2036–2042 (2022).

48. Pryor, J. M. et al. Enabling one-pot Golden Gate assemblies of unprecedented complexity using data-optimized assembly design. PLoS ONE 15, e0238592 (2020).

49. Potapov, V. et al. Comprehensive Profiling of Four Base Overhang Ligation Fidelity by T4 DNA Ligase and Application to DNA Assembly. ACS Synth. Biol. 7, 2665–2674 (2018).

50. Wick, S. et al. PERSIA for Direct Fluorescence Measurements of Transcription, Translation, and Enzyme Activity in Cell-Free Systems. ACS Synth. Biol. 8, 1010–1025 (2019).

51. Willi, J. A., Karim, A. S. & Jewett, M. C. Cell-Free Translation Quantification via a Fluorescent Minihelix. ACS Synth. Biol. 13, 2253–2259 (2024).

52. Kahn, T. W., Beachy, R. N. & Falk, M. M. Cell-free expression of a GFP fusion protein allows quantitation in vitro and in vivo. Current Biology 7, R207–R208 (1997).

53. Kouba, P. et al. Machine Learning-Guided Protein Engineering. ACS Catal. 13, 13863–13895 (2023).

54. Nayab, A., Moududee, S. A., Shi, Y., Jiang, Y. & Gong, Q. Crystal Structure of Urate Oxidase from Bacillus Subtilis 168. Crystallogr. Rep. 64, 1126–1133 (2019).

55. Galloway Lab. Oligo annealing for ligation cloning. Galloway Lab Protocols https://gallowaylabmit.github.io/protocols/en/latest/protocols/cloning/oligo_annealing.html (2023).

56. Hejazi, S., Godin, R., Jurasic, V. & Reuel, N. F. Single-Walled Carbon Nanotube Probes for Protease Characterization Directly in Cell-Free Expression Reactions. Anal. Chem. 97, 10745–10754 (2025).

