## Supporting Information for "Rapid Cell-Free Combinatorial Mutagenesis Workflow Using Small Oligos Suitable for High-Iteration, Active Learning-Guided Protein Engineering"

### Table of Contents

**Supplementary Tables** **S-2**

*Table S1. Sequences for Full-Length Gene Fragments Used in This Study* S-2

*Table S2. Sequences for Partial Gene Fragments Used in This Study* S-4

*Table S3. Sequences for Primers Used in This Study* S-5

*Table S4. Sequences for Annealed Oligos Used in This Study* S-5

**Supplementary Methods** **S-6**

**Supplementary Data** **S-8**

*Figure S1. Complete Sanger Sequencing Data* S-8

*Figure S2. SfGFP Calibration Curve* S-9

*Figure S3. Split GFP Calibration Curve* S-9

*Figure S4. HaloTag Normalization Data* S-10

### Supplementary Tables

**Table S1. Sequences for Full-Length Gene Fragments Used in This Study.**

| Name | Primers | Sequence |
|---|---|---|
| Sequence Info | | Primer-T7 Promoter-Ribosome Binding Site-Start Codon-Golden Gate Site-Mutation Site-Strep-II Tag-Stop Codon-T7 Terminator |
| sfGFP LET | Forward: M13-Forward<br><br>Reverse: M13-Reverse | GTAAAACGACGGCCAGTAGCGCTATTAAAGCTTCGAAATTAAT<br>ACGACTCACTATAGGGAGACCACAACGGTTTCCCTCTAGAAA<br>TAATTTTGTTTAACTTTAAGAAGGAGATATACATATGAGCAAAG<br>GTGAAGAACTGTTTACCGGCGTTGTGCCGATTCTGGTGGAAC<br>TGGATGGCGATGTGAACGGTCACAAATTCAGCGTGCGTGGT<br>GAAGGTGAAGGCGATGCCACGATTGGCAAACTGACGCTGAA<br>ATTTATCTGCACCACCGGCAAACTGCCGGTGCCGTGGCCGA<br>CGCTGGTGACCACCCTGACCTATGGCGTTCAGTGTTTTAGTC<br>GCTATCCGGATCACATGAAACGTCACGATTTCTTTAAATCTGC<br>AATGCCGGAAGGCTATGTGCAGGAACGTACGATTAGCTTTAA<br>AGATGATGGCAAATATAAAACGCGCGCCGTTGTGAAATTTGAA<br>GGCGATACCCTGGTGAACCGCATTGAACTGAAAGGCACGGA<br>TTTTAAAGAAGATGGCAATATCCTGGGCCATAAACTGGAATAC<br>AACTTTAATAGCCATAATGTTTATATTACGGCGGATAAACAGAA<br>AAATGGCATCAAAGCGAATTTTACCGTTCGCCATAACGTTGAA<br>GATGGCAGTGTGCAGCTGGCAGATCATTATCAGCAGAATACC<br>CCGATTGGTGATGGTCCGGTGCTGCTGCCGGATAATCATTAT<br>CTGAGCACGCAGACCGTTCTGTCTAAAGATCCGAACGAAAAA<br>CGGGACCACATGGTTCTGCACGAATATGTGAATGCGGCAGGT<br>ATTACGTGGAGCCATCCGCAGTTCGAAAAATAATAAGTCGAC<br>CGGCTGCTAACAAAGCCCGAAAGGAAGCTGAGTTGGCTGCT<br>GCCACCGCTGAGCAATAACTAGCATAACCCCTTGGGGCCTCT<br>AAACGGGTCTTGAGGGGTTTTTTGCTGAAAGCGAGACTAAGC<br>TTTAAACTTCGGGTCATAGCTGTTTCCTG |
| Uricase-SplitGFP LET | Forward: M13-Forward<br><br>Reverse: M13-Reverse | GTAAAACGACGGCCAGTAGCGCTATTAAAGCTTCGAAATTAAT<br>ACGACTCACTATAGGGAGAGCACAACGGTTTCCCTCTAGAAA<br>TAATTTTGTTTAACTTTAAGAAGGAGATATACATATGAAACGGA<br>CCATGAGTTACGGGAAGGGCAACGTGTTCGCCTACCGGACG<br>TATCTCAAGCCTCTCACCGGGGTGAAGCAGATCCCAGAATCA<br>AGCTTCGCCGGGCGGGATAATACTGTTGTTGGTGTTGATGTG<br>ACATGCGAAATTGGCGGTGAAGCCTTCTTACCAAGTTTCACA<br>GATGGTGATAACACTCTGGTCGTCGCAACGGACTCTATGAAG<br>AATTTTATTCAGCGGCACTTGGCAAGCTATGAAGGGACTACAA<br>CAGAGGGTTTCCTCCATTATGTGGCGCACCGGTTTTTAGATAC<br>GTACTCACACATGGACACAATCACCCTGACTGGGGAGGACAT<br>TCCGTTTGAGGCCATGCCAGCTTACGAAGAGAAGGAATTGTC<br>GACGTCTCGTTTGGTGTTTCGTCGCTCCCGTAATGAGCGTTC<br>TCGTTCAGTACTTAAAGCAGAGCGGTCAGGTAACACTATCACA<br>ATCACTGAGCAGTACTCGGAGATCATGGACTTACAACTTGTAA<br>AGGTTAGTGGCAATAGCTTTGTGGGCTTCATTCGTGATGAGTA<br>CACCACATTACCAGAGGATGGCAACCGGCCGCTGTTCGTCTA<br>CCTGAATATTTCCTGGCAGTACGAAAATACGAATGATTCTTAC<br>GCTAGCGATCCTGCCCGCTATGTAGCTGCCGAACAAGTTCGT<br>GACCTTGCGAGTACTGTTTTCCATGAGCTTGAGACGCCTTCC<br>ATTCAGAACTTGATCTATCATATCGGCTGCCGTATCCTCGCCC<br>GTTTCCCACAGCTCACGGACGTCAGCTTTCAATCGCAAAATC<br>ATACGTGGGACACCGTGGTCGAGGAAATCCCTGGCTCGAAG<br>GGGAAGGTATATACTGAACCGCGGCCGCCGTACGGTTTTCAA<br>CATTTCACTGTTACTCGGGAAGACGCTGAGAAGGAAAAGCAA<br>AAGGCAGCCGAGAAATGTCGGTCGTTGAAGGCACTGATTGG<br>CAGCGATGGCGGCAGCGGCGGCGGCAGCACCAGCCGCGAT |

| | | |
|---|---|---|
| | | CATATGGTGCTGCATGAATATGTGAACGCGGCGGGCATTACC<br>AGCGCGTGGAGCCATCCGCAGTTTGAAAAATAATAAGTCGAC<br>CGGCTGCTAACAAAGCCCGAAAGGAAGCTGAGTTGGCTGCT<br>GCCACCGCTGAGCAATAACTAGCATAACCCCTTGGGGCCTCT<br>AAACGGGTCTTGAGGGGTTTTTTGCTGAAAGCGAGACTAAGC<br>TTTAAACTTCGGGTCATAGCTGTTTCCTG |
| Uricase-HaloTag LET | Forward: M13-Forward<br><br>Reverse: M13-Reverse | GTAAAACGACGGCCAGTAGCGCTATTAAAGCTTCGAAATTAAT<br>ACGACTCACTATAGGGAGACCACAACGGTTTCCCTCTAGAAA<br>TAATTTTGTTTAACTTTAAGAAGGAGATATACATATGAAACGGA<br>CCATGAGTTACGGGAAGGGCAACGTGTTCGCCTACCGGACG<br>TATCTCAAGCCTCTCACCGGGGTGAAGCAGATCCCAGAATCA<br>AGCTTCGCCGGGCGGGATAATACTGTTGTTGGTGTTGATGTG<br>ACATGCGAAATTGGCGGTGAAGCCTTCTTACCAAGTTTCACA<br>GATGGTGATAACACTCTGGTCGTCGCAACGGACTCTATGAAG<br>AATTTTATTCAGCGGCACTTGGCAAGCTATGAAGGGACTACAA<br>CAGAGGGTTTCCTCCATTATGTGGCGCACCGGTTTTTAGATAC<br>GTACTCACACATGGACACAATCACCCTGACTGGGGAGGACAT<br>TCCGTTTGAGGCCATGCCAGCTTACGAAGAGAAGGAATTGTC<br>GACGTCTCGTTTGGTGTTTCGTCGCTCCCGTAATGAGCGTTC<br>TCGTTCAGTACTTAAAGCAGAGCGGTCAGGTAACACTATCACA<br>ATCACTGAGCAGTACTCGGAGATCATGGACTTACAACTTGTAA<br>AGGTTAGTGGCAATAGCTTTGTGGGCTTCATTCGTGATGAGTA<br>CACCACATTACCAGAGGATGGCAACCGGCCGCTGTTCGTCTA<br>CCTGAATATTTCCTGGCAGTACGAAAATACGAATGATTCTTAC<br>GCTAGCGATCCTGCCCGCTATGTAGCTGCCGAACAAGTTCGT<br>GACCTTGCGAGTACTGTTTTCCATGAGCTTGAGACGCCTTCC<br>ATTCAGAACTTGATCTATCATATCGGCTGCCGTATCCTCGCCC<br>GTTTCCCACAGCTCACGGACGTCAGCTTTCAATCGCAAAATC<br>ATACGTGGGACACCGTGGTCGAGGAAATCCCTGGCTCGAAG<br>GGGAAGGTATATACTGAACCGCGGCCGCCGTACGGTTTTCAA<br>CATTTCACTGTTACTCGGGAAGACGCTGAGAAGGAAAAGCAA<br>AAGGCAGCCGAGAAATGTCGGTCGTTGAAGGCATTGGAACC<br>TACAACAGAAGACCTGTACTTCCAATCGGATAACGATGGGTCA<br>GAAATTGGTACCGGTTTTCCGTTCGATCCGCATTACGTAGAG<br>GTTTTGGGGGAACGTATGCACTACGTTGACGTTGGCCCTCGG<br>GATGGGACCCCAGTACTCTTTTTGCACGGCAACCCTACTAGC<br>TCATATGTTTGGCGGAATATTATTCCACATGTAGCGCCGACCC<br>ACCGGTGTATCGCACCGGACCTGATTGGGATGGGCAAGTCG<br>GATAAGCCAGACCTTGGGTACTTTTTCGACGATCACGTACGC<br>TTTATGGACGCTTTCATTGAGGCCTTAGGCCTGGAAGAGGTA<br>GTTTTAGTAATCCATGACTGGGGGTCAGCCTTGGGCTTTCAC<br>TGGGCTAAACGGAATCCAGAGCGTGTGAAGGGTATTGCATTC<br>ATGGAATTCATCCGCCCGATCCCGACATGGGATGAGTGGCCA<br>GAGTTTGCACGTGAAACATTTCAAGCTTTTCGGACCACAGAC<br>GTTGGCCGCAAGTTGATCATCGATCAAAACGTGTTCATCGAA<br>GGGACTCTCCCAATGGGTGTCGTCCGCCCGTTGACTGAGGT<br>CGAGATGGACCACTATCGGGAGCCTTTCTTAAACCCAGTAGA<br>TCGGGAACCGTTGTGGCGGTTCCCGAACGAATTGCCAATTG<br>CAGGCGAGCCAGCAAACATCGTAGCCCTCGTGGAGGAGTAC<br>ATGGACTGGCTTCACCAGTCGCCTGTGCCTAAATTACTGTTCT<br>GGGGGACTCCTGGTGTCCTTATCCCTCCTGCGGAGGCTGCC<br>CGTTTAGCGAAGTCGTTGCCAAATTGCAAGGCAGTGGATATC<br>GGCCCGGGTCTTAACCTTTTGCAAGAAGATAACCCGGACTTA<br>ATTGGTTCTGAAATTGCTCGTTGGCTCAGTACCCTCGAAATTA<br>GTGGCCATCATCACCATCACCACTAATAAGTCGACCGGCTGC<br>TAACAAAGCCCGAAAGGAAGCTGAGTTGGCTGCTGCCACCG<br>CTGAGCAATAACTAGCATAACCCCTTGGGGCCTCTAAACGGG<br>TCTTGAGGGGTTTTTTGCTGAAAGCGAGACTAAGCTTTAAACT<br>TCGGGTCATAGCTGTTTCCTG |

**Table S2. Sequences for Partial Gene Fragments Used in This Study.**

| Name | Primers | Sequence |
|---|---|---|
| SfGFP-Left | Forward: M13-Forward<br><br>Reverse: SfGFP-Left-R | GTAAAACGACGGCCAGTAGCGCTATTAAAGCTTCGAAATTAAT<br>ACGACTCACTATAGGGAGACCACAACGGTTTCCCTCTAGAAA<br>TAATTTTGTTTAACTTTAAGAAGGAGATATACATATGAGCAAAG<br>GTGAAGAACTGTTTACCGGCGTTGTGCCGATTCTGGTGGAAC<br>TGGATGGCGATGTGAACGGTCACAAATTCAGCGTGCGTGGT<br>GAAGGTGAAGGCGATGCCACGATTGGCAAACTGACGCTGAA<br>ATTTATCTGCACCACCGGCAAACTGCCGGTGCCGTGGCCGA<br>CGCTGGTGACCACCCTGACCT |
| SfGFP-Right | Forward: SfGFP-Right-F<br><br>Reverse: M13-Reverse | TTTCTTTAAATCTGCAATGCCGGAAGGCTATGTGCAGGAACGT<br>ACGATTAGCTTTAAAGATGATGGCAAATATAAAACGCGCGCCG<br>TTGTGAAATTTGAAGGCGATACCCTGGTGAACCGCATTGAAC<br>TGAAAGGCACGGATTTTAAAGAAGATGGCAATATCCTGGGCC<br>ATAAACTGGAATACAACTTTAATAGCCATAATGTTTATATTACGG<br>CGGATAAACAGAAAAATGGCATCAAAGCGAATTTTACCGTTCG<br>CCATAACGTTGAAGATGGCAGTGTGCAGCTGGCAGATCATTA<br>TCAGCAGAATACCCCGATTGGTGATGGTCCGGTGCTGCTGCC<br>GGATAATCATTATCTGAGCACGCAGACCGTTCTGTCTAAAGAT<br>CCGAACGAAAAACGGGACCACATGGTTCTGCACGAATATGTG<br>AATGCGGCAGGTATTACGTGGAGCCATCCGCAGTTCGAAAAA<br>TAATAAGTCGACCGGCTGCTAACAAAGCCCGAAAGGAAGCTG<br>AGTTGGCTGCTGCCACCGCTGAGCAATAACTAGCATAACCCC<br>TTGGGGCCTCTAAACGGGTCTTGAGGGGTTTTTTGCTGAAAG<br>CGAGACTAAGCTTTAAACTTCGGGTCATAGCTGTTTCCTG |
| Uricase-Left | Forward: M13-Forward<br><br>Reverse: Uricase-Reverse | GTAAAACGACGGCCAGTAGCGCTATTAAAGCTTCGAAATTAAT<br>ACGACTCACTATAGGGAGAGCACAACGGTTTCCCTCTAGAAA<br>TAATTTTGTTTAACTTTAAGAAGGAGATATACATATGAAACGGA<br>CCATGAGTTACGGGAAGGGCAACGTGTTCGCCTACCGGACG<br>TATCTCAAGCCTCTCACCGGGGTGAAGCAGATCCCAGAATCA<br>AGCTTCGCCGGGCGGGAGACCAAGATAGCGT |
| Uricase-Spacer | Forward: Uricase-Forward<br><br>Reverse: Uricase-Reverse | GTAAGCCTACGGTCTCTACATGCGAAATTGGCGGTGAAGCCT<br>TCTTACCAAGTTTCACAGATGGTGATAACACTCTGGTCGTCGC<br>AACGGACTCTATGAAGAATTTTATTCAGCGGCACTTGGCAAGC<br>TATGAAGGGACTACAACAGAGGGTTTCCTCCATTATGTGGCG<br>CACCGGTTTTTAGATACGTACTCACACATGGACACAATCACCC<br>TGACTGGGGAGGACATTCCGTTTGAGGCCATGCCAGCTTAC<br>GAAGAGAAGGAATTGTCGACGTCTCGTTTGGTGTTTCGTCGC<br>TCCCGTAATGAGCGTTCTCGTTCAGTACTTAAAGCAGAGCGG<br>TCAGGTAACACTATCACAATCACTGAGCAGTACTCGGAGATCA<br>TGGACTTACAACTTGTAAAGGTTAGTGGCAATAGCTTTGTGGG<br>CTTCATTCGTGATGAGTACACCACATTACCAGAGGATGGCAAC<br>CGGCCGCTGTTCGTCTACCTGAATATTTCCTGGCAGTACGAA<br>AATACGAATGATTCTTACGCTAGCGATCCTGCCCGCTATGTAG<br>CTGCCGAACAAGTTCGTGACCTTGCGAGTACTGTTTTCCATG<br>AGCTTGAGACGCCTTCCATTCAGAACTTGATCTATCATATCGG<br>CTGCCGTATCCTCGCCCGTTTCCCACAGCTCACGGACGTCA<br>GCGGAGACCAAGATAGCGT |
| Uricase-Right | Forward: Uricase-Forward<br><br>Reverse: M13-Reverse | GTAAGCCTACGGTCTCTTCACTGTTACTCGGGAAGACGCTGA<br>GAAGGAAAAGCAAAAGGCAGCCGAGAAATGTCGGTCGTTGA<br>AGGCACTGATTGGCAGCGATGGCGGCAGCGGCGGCGGCAG<br>CACCAGCCGCGATCATATGGTGCTGCATGAATATGTGAACGC<br>GGCGGGCATTACCAGCGCGTGGAGCCATCCGCAGTTTGAAA<br>AATAATAAGTCGACCGGCTGCTAACAAAGCCCGAAAGGAAGC<br>TGAGTTGGCTGCTGCCACCGCTGAGCAATAACTAGCATAACC<br>CCTTGGGGCCTCTAAACGGGTCTTGAGGGGTTTTTTGCTGAA<br>AGCGAGACTAAGCTTTAAACTTCGGGTCATAGCTGTTTCCTG |
| sfGFP-10-Left | Forward: M13-Forward<br><br>Reverse: SfGFP-10-Left-R | GTAAAACGACGGCCAGTAGCGCTATTAAAGCTTCGAAATTAAT<br>ACGACTCACTATAGGGAGACCACAACGGTTTCCCTCTAGAAA<br>TAATTTTGTTTAACTTTAAGAAGGAGATATACATATGAGCAAAG<br>GTGAAGAACTGTTTACCGGCGTTGTGCCGATTCTGGTGGAAC |

| | | |
|---|---|---|
| | | TGGATGGCGATGTGAACGGTCACAAATTCAGCGTGCGTGGT GAAGGTGAAGGCGATGCCACGATTGGCAAACTGACGCTGAA ATTTATCTGCACCACCGGCAAACTG |
| sfGFP-10-Right | Forward: SfGFP-10-Right-F<br><br>Reverse: M13-Reverse | CGATACCCTGGTGAACCGCATTGAACTGAAAGGCACGGATTT TAAAGAAGATGGCAATATCCTGGGCCATAAACTGGAATACAAC TTTAATAGCCATAATGTTTATATTACGGCGGATAAACAGAAAAAT GGCATCAAAGCGAATTTTACCGTTCGCCATAACGTTGAAGATG GCAGTGTGCAGCTGGCAGATCATTATCAGCAGAATACCCCGA TTGGTGATGGTCCGGTGCTGCTGCCGGATAATCATTATCTGA GCACGCAGACCGTTCTGTCTAAAGATCCGAACGAAAAACGG GACCACATGGTTCTGCACGAATATGTGAATGCGGCAGGTATTA CGTGGAGCCATCCGCAGTTCGAAAAATAATAAGTCGACCGGC TGCTAACAAAGCCCGAAAGGAAGCTGAGTTGGCTGCTGCCA CCGCTGAGCAATAACTAGCATAACCCCTTGGGGCCTCTAAAC GGGTCTTGAGGGGTTTTTTGCTGAAAGCGAGACTAAGCTTTA AACTTCGGGTCATAGCTGTTTCCTG |

**Table S3. Sequences for Primers Used in This Study**

| Name | Sequence |
|---|---|
| M13-Forward | GTAAAACGACGGCCAGT |
| M13-Reverse | CAGGAAACAGCTATGAC |
| SfGFP-Left-R | GGCTACCGTCTCAGGTCAGGGTGGTCAC |
| SfGFP-Right-F | GGCTACCGTCTCTTTCTTTAAATCTGCAATGCC |
| SfGFP-10-Left-R | GGCTACCGTCTCAGTTTGCCGGTGGTG |
| SfGFP-10-Right-F | GGCTACCGTCTCCGATACCCTGGTGAAC |
| Uricase-Forward | GTAAGCCTACGGTCTCT |
| Uricase-Reverse | ACGCTATCTTGGTCTCC |
| UricaseHalo-Frag1-R* | GGCTACGGTCTCGTCACATCAACACCAACAACA |
| UricaseHalo-Frag2-F | GGCTACGGTCTCTGTGACATGCGAAATTGGC |
| UricaseHalo-Frag2-R | GGCTACGGTCTCTTTGCGATTGAAAGCTGACG |
| UricaseHalo-Frag3-F | GGCTACGGTCTCCGCAAAATCATACGTGGGAC |
| UricaseHalo-Frag3-R | GGCTACGGTCTCACCTTCCCCTTCGAGC |
| UricaseHalo-Frag4-F* | GGCTACGGTCTCGAAGGTATATACTGAACCGC |
| UricaseHalo-Frag1-D44V-R | GGCTACGGTCTCGTCACAACAACACCAACAACA |
| UricaseHalo-Frag2-Q268R-R | GGCTACGGTCTCTTTGCGATCGAAAGCTGACG |
| UricaseHalo-Frag3-K285Q-R | GGCTACGGTCTCACCTTCCCCTGCGAGCC |

*The UricaseHalo-Frag1-R and UricaseHalo-Frag1-F primers are paired with the M13-Forward and M13-Reverse primers. The mutant primers (e.g. UricaseHalo-Frag1-D44V-R) are paired with the forward primer for that fragment.

**Table S4. Sequences for Annealed Oligos Used in This Study**

| Name | Sequence |
|---|---|
| *sfGFP Annealed Oligos* | |
| Anneal1 (10-Anneal3) | F: GACCTATGGCGTTCAGTGT<br>R: TAAAACACTGAACGCCATA |
| Anneal2 (10-Anneal4) | F: TTTAGTCGCTATCCGGAT<br>R: TGTGATCCGGATAGCGAC |
| Anneal3 (10-Anneal5) | F: CACATGAAACGTCACGAT<br>R: AGAAATCGTGACGTTTCA |
| 10-Anneal1 | F: AAACTGCCGGTGCCGTGG<br>R: TCGGCCACGGCACCGGCA |
| 10-Anneal2 | F: CCGACGCTGGTGACCACCCT<br>R: GGTCAGGGTGGTCACCAGCG |
| 10-Anneal6 | F: TTCTTTAAATCTGCAATGCCG<br>R: CTTCCGGCATTGCAGATTTAA |
| 10-Anneal7 | F: GAAGGCTATGTGCAGGAA<br>R: TACGTTCCTGCACATAGC |

| | |
|---|---|
| 10-Anneal8 | F: CGTACGATTAGCTTTAAAGATG<br>R: CCATCATCTTTAAAGCTAATCG |
| 10-Anneal9 | F: ATGGCAAATATAAAACGCGC<br>R: ATGGCAAATATAAAACGCGC |
| 10-Anneal10 | F: GCCGTTGTGAAATTTGAAGGC<br>R: TATCGCCTTCAAATTTCACAA |
| Anneal1-Y66H (10-Anneal3-Y66H) | F: GACCCATGGCGTTCAGTGT<br>R: TAAAACACTGAACGCCATG |
| Anneal1-Y66W (10-Anneal3-Y66W) | F: GACCTGGGGCGTTCAGTGT<br>R: TAAAACACTGAACGCCCCA |
| *Uricase Annealed Oligos* | |
| Anneal1 | F: GGCGGGATAATACTGTTGTTGGTGTTGATGTG<br>R: ATGTCACATCAACACCAACAACAGTATTATCC |
| Anneal2 | F: CAGCTTTCAATCGCAAAATCATA<br>R: CACGTATGATTTTGCGATTGAAA |
| Anneal3 | F: CGTGGGACACCGTGGTCGAGGAA<br>R: GGATTTCCTCGACCACGGTGTCC |
| Anneal4 | F: ATCCCTGGCTCGAAGGGGAAGGTATA<br>R: AGTATATACCTTCCCCTTCGAGCCAG |
| Anneal5 | F: TACTGAACCGCGGCCGCCGTACGGTTTTCAACATT<br>R: GTGAAATGTTGAAAACCGTACGGCGGCCGCGGTTC |
| Anneal1-D44V | F: GGCGGGATAATACTGTTGTTGGTGTTGTTGTG<br>R: ATGTCACAACAACACCAACAACAGTATTATCC |
| Anneal2-Q268R | F: CAGCTTTCGATCGCAAAATCATA<br>R: CACGTATGATTTTGCGATCGAAA |
| Anneal3-E279K | F: CGTGGGACACCGTGGTCAAGGAA<br>R: GGATTTCCTTGACCACGGTGTCC |
| Anneal4-K285Q | F: ATCCCTGGCTCGCAGGGGAAGGTATA<br>R: AGTATATACCTTCCCCTGCGAGCCAG |
| Anneal5-H300R | F: TACTGAACCGCGGCCGCCGTACGGTTTTCAACGTT<br>R: GTGAAACGTTGAAAACCGTACGGCGGCCGCGGTTC |

**Supplementary Methods**

HaloTag Mutant Preparation:

Uricase mutants tagged with the HaloTag® protein were prepared using Golden Gate assembly from larger, mutagenic gene fragments prepared using site-directed mutagenesis, not small annealed-oligos like in the main paper. The exact DNA sequence information is provided above. Specifically, the sequence was divided into four regions containing the positions for D44, Q268, K285, and the HaloTag®. Both the wild-type fragments and the mutated fragments (for D44V, Q268R, and K285Q) were PCR-amplified from the full-length gBlock LET template using the primers specified, purified, and quantified as described in the main methods. Once purified, Golden Gate reactions were setup for the desired mutants using 1 µL of the required fragments,

2 μL of 10X T4 DNA Ligase Buffer, 1 μL of BsaI-HF-v2 enzyme mix, and 13 μL of nuclease-free water using the same parameters as before. The resulting mixture was again amplified, purified, and quantified for use in cell-free expression reactions. However, the elongation time during the PCR was changed to from 30 s to 1 min.

HaloTag Assay:

The uricase activity was determined using the Amplex™ Red Uric Acid/Uricase Assay Kit (#A22181, ThermoFisher). Specifically, purified DNA encoding the protein variants was expressed with cell-free expression at a 5 nM load overnight at 30°C. After, incubation, 2 μL of cell-free expression product was diluted with 48 μL of assay buffer and subsequently mixed with 50 μL of working solution according to kit instructions (working solution was made with 50% more 5 mM uric acid solution than recommended) in a black, flat transparent bottom 384-well plate, the plate was sealed with an Axygen PCR-SP plate film, and the fluorescence was measured (excitation/emission: 540/590, bandwith of 25 and 20, respectively, and a gain of 80) every minute for 1 hour at 37°C with continuous shaking at 240 rpm. The data for the first 15 min was fit to a linear regression curve after background subtraction with a no DNA control and the slope was taken as the activity. For the HaloTag® expression measurement, 13 μL of cell-free expression product was mixed with 5 μL of a 10 μM Janelia Fluor® 646 HaloTag® ligand solution (20% DMSO in 50 mM HEPES buffer at pH = 7) in a black, flat transparent bottom 384-well plate, the plate was sealed with an Axygen PCR-SP plate film, and the fluorescence was measured (excitation/emission: 620/680, bandwith of 25 and 30, respectively, and a gain of 120) every minute for 1 hour at 37°C with continuous shaking at 240 rpm. The reading at 40 min was used as the signal after background subtraction with a no DNA control. The normalized activity was calculated by dividing the uricase activity by the HaloTag® signal.

### Supplementary Data

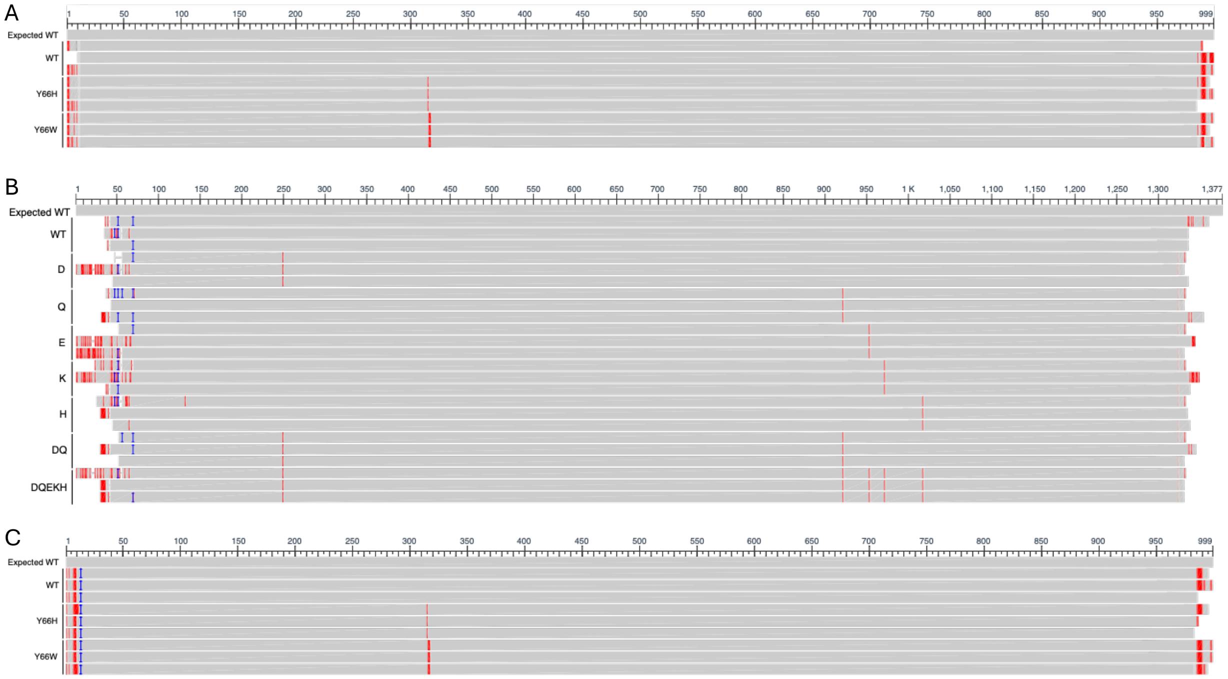


**Figure S1. Complete Sanger Sequencing Data.** Multiple sequence alignment of Sanger sequencing data for the (A) 3-oligo sfGFP assembly, (B) 5-oligo uricase assembly, and (C) 10-oligo sfGFP assembly. Data shows substitutions at the target mutations sites for multiple mutants and n = 3 independent assemblies.

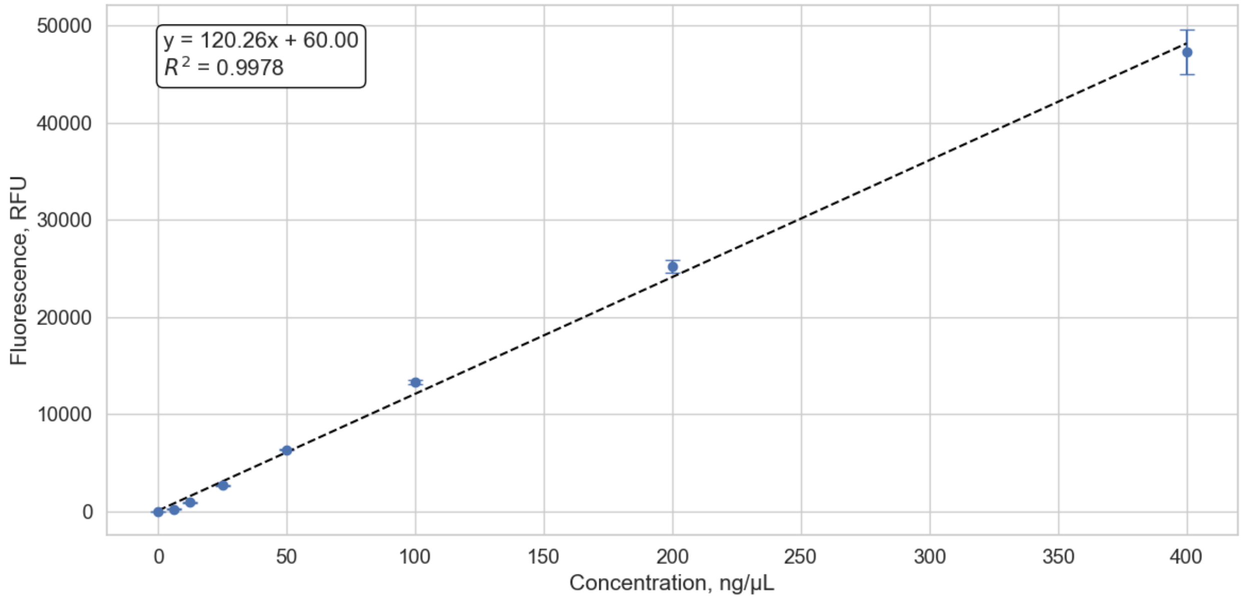


**Figure S2. SfGFP Calibration Curve.** Calibration curve was constructed using purified sfGFP protein expressed from the full-length PCR-amplified IDT gBlock. Expression and purification is as detailed in the main methods. Data represents the average of n = 3 technical replicates. Error bars represent the standard deviation from the mean.

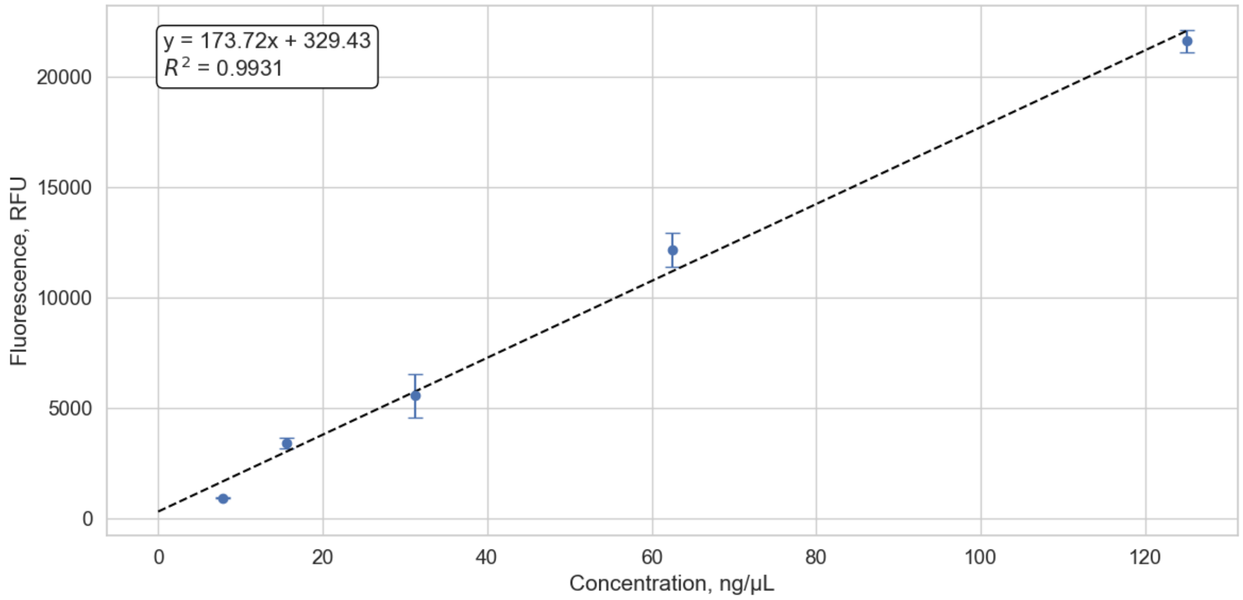


**Figure S3. Split GFP Calibration Curve.** Calibration curve was constructed using purified GFP11-tagged Uricase protein expressed from the full-length PCR-amplified IDT gBlock. Expression and purification is as detailed in the main methods. Data represents the average of n = 3 technical replicates. Error bars represent the standard deviation from the mean.

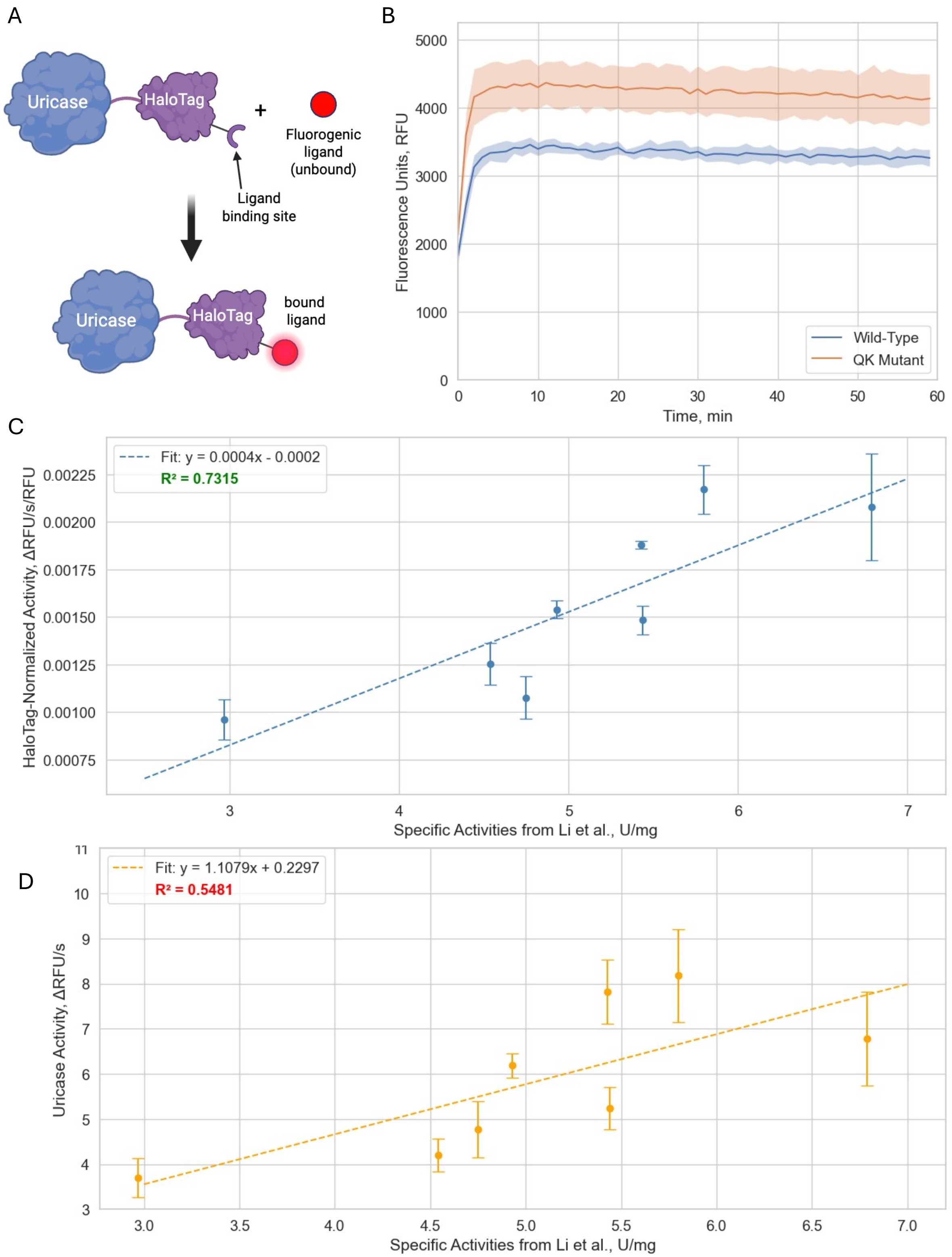


**Figure S4. HaloTag Normalization Data.** (A) Overview of HaloTag® quantification strategy. It relies on activation of a fluorogenic dye (Janelia Fluor® 646 HaloTag® ligand) upon binding to the HaloTag® to produce a fluorescence signal proportional to protein concentration so that the concentration can be calculated directly in lysate. Like the GFP11-tag and split GFP system, this allows protein activities calculated with lysate-based screens to be normalized to protein amount and minimize biases from expression level differences. (B) Kinetic curve showing rapid signal stabilization after just 10 min for two different uricase mutant proteins. (C,D) Comparison of correlation of normalized and non-normalized activities to the specific activity values previously reported in Li et al. The following mutants were analyzed: WT, D, Q, K, DQ, DK, QK, and DQK. All data represents the average of n = 3 independent replicates. Error bars represent the standard deviation from the mean.
