## Supporting FIle 1 for "Rapid Cell-Free Combinatorial Mutagenesis Workflow Using Small Oligos Suitable for High-Iteration, Active Learning-Guided Protein Engineering": TestGoldenGateOT2.py

```
from opentrons import protocol_api
import os, math, io, string, pandas as pd

# metadata
metadata = {
    "protocolName": "GGA Pipeline",
    "author": "Ryan Godin",
    "description": "Simple protocol to test the GGA workflow",
}

# requirements
requirements = {"robotType": "OT-2", "apiLevel": "2.15"}

# Data
seqFileCSV = '''Well ID,Sequence
A1,"FragMix, Anneal2, Anneal6m-Q268R, Anneal7m-E279K, Anneal8m-K285Q, Anneal9m-H300R"
B1,"FragMix, Anneal2m-D44V, Anneal6m-Q268R, Anneal7m-E279K, Anneal8, Anneal9m-H300R"
C1,"FragMix, Anneal2m-D44V, Anneal6m-Q268R, Anneal7, Anneal8m-K285Q, Anneal9m-H300R"
D1,"FragMix, Anneal2m-D44V, Anneal6m-Q268R, Anneal7m-E279K, Anneal8m-K285Q, Anneal9"
E1,"FragMix, Anneal2, Anneal6, Anneal8, Anneal9"
F1,"FragMix, Anneal2m-D44V, Anneal6m-Q268R, Anneal7m-E279K, Anneal8, Anneal9"
G1,"FragMix, Anneal2m-D44V, Anneal6m-Q268R, Anneal7, Anneal8, Anneal9"
H1,"FragMix, Anneal2m-D44V, Anneal6m-Q268R, Anneal7m-E279K, Anneal8m-K285Q, Anneal9"
I1,"FragMix, Anneal2m-D44V, Anneal6m-Q268R, Anneal7m-E279K, Anneal8m-K285Q, Anneal9"
J1,"FragMix, Anneal2, Anneal6, Anneal7, Anneal8, Anneal9m-H300R"
K1,"FragMix, Anneal2m-D44V, Anneal6, Anneal7m-E279K, Anneal8m-K285Q, Anneal9m-H300R"
L1,"FragMix, Anneal2m-D44V, Anneal6, Anneal7, Anneal8, Anneal9"
M1,"FragMix, Anneal2m-D44V, Anneal6m-Q268R, Anneal7m-E279K, Anneal8m-K285Q, Anneal9m-H300R"
N1,"FragMix, Anneal2m-D44V, Anneal6, Anneal7, Anneal8, Anneal9"
O1,"FragMix, Anneal2, Anneal6, Anneal7, Anneal8m-K285Q, Anneal9"
P1,"FragMix, Anneal2, Anneal6, Anneal7m-E279K, Anneal8, Anneal9"
A2,"FragMix, Anneal2m-D44V, Anneal6m-Q268R, Anneal7m-E279K, Anneal8m-K285Q, Anneal9"
B2,"FragMix, Anneal2, Anneal6, Anneal7, Anneal8m-K285Q, Anneal9"
C2,"FragMix, Anneal2, Anneal6, Anneal7, Anneal8m-K285Q, Anneal9"
D2,"FragMix, Anneal2m-D44V, Anneal6m-Q268R, Anneal7, Anneal8, Anneal9"
E2,"FragMix, Anneal2, Anneal6m-Q268R, Anneal7, Anneal8, Anneal9"
F2,"FragMix, Anneal2, Anneal6, Anneal8, Anneal9"
G2,"FragMix, Anneal2m-D44V, Anneal6m-Q268R, Anneal7, Anneal8m-K285Q, Anneal9m-H300R"
H2,"water1, water2, water3, water4, water5"
I2,"FragMix, Anneal2, Anneal6, Anneal7, Anneal8, Anneal9m-H300R"
J2,"FragMix, Anneal2, Anneal6m-Q268R, Anneal7m-E279K, Anneal8m-K285Q, Anneal9m-H300R"
K2,"FragMix, Anneal2, Anneal6, Anneal7, Anneal8, Anneal9m-H300R"
L2,"FragMix, Anneal2, Anneal6, Anneal8, Anneal9"
M2,"FragMix, Anneal2m-D44V, Anneal6, Anneal7, Anneal8, Anneal9"
N2,"FragMix, Anneal2, Anneal6, Anneal7m-E279K, Anneal8, Anneal9"
O2,"FragMix, Anneal2m-D44V, Anneal6m-Q268R, Anneal7, Anneal8, Anneal9"
P2,"FragMix, Anneal2m-D44V, Anneal6m-Q268R, Anneal7m-E279K, Anneal8, Anneal9"
A3,"FragMix, Anneal2, Anneal6, Anneal8, Anneal9"
B3,"FragMix, Anneal2m-D44V, Anneal6m-Q268R, Anneal7, Anneal8m-K285Q, Anneal9m-H300R"
C3,"FragMix, Anneal2, Anneal6, Anneal7, Anneal8m-K285Q, Anneal9"
D3,"FragMix, Anneal2, Anneal6m-Q268R, Anneal7, Anneal8, Anneal9"
E3,"FragMix, Anneal2, Anneal6, Anneal7m-E279K, Anneal8, Anneal9"
F3,"water1, water2, water3, water4, water5"
G3,"FragMix, Anneal2, Anneal6m-Q268R, Anneal7, Anneal8, Anneal9"
H3,"FragMix, Anneal2, Anneal6, Anneal7, Anneal8m-K285Q, Anneal9"
I3,"FragMix, Anneal2, Anneal6m-Q268R, Anneal7m-E279K, Anneal8m-K285Q, Anneal9m-H300R"
J3,"FragMix, Anneal2m-D44V, Anneal6m-Q268R, Anneal7, Anneal8, Anneal9"
K3,"FragMix, Anneal2m-D44V, Anneal6, Anneal7m-E279K, Anneal8m-K285Q, Anneal9m-H300R"
L3,"FragMix, Anneal2m-D44V, Anneal6m-Q268R, Anneal7m-E279K, Anneal8, Anneal9m-H300R"
M3,"FragMix, Anneal2m-D44V, Anneal6m-Q268R, Anneal7, Anneal8m-K285Q, Anneal9m-H300R"
N3,"water1, water2, water3, water4, water5"
O3,"FragMix, Anneal2m-D44V, Anneal6m-Q268R, Anneal7m-E279K, Anneal8m-K285Q, Anneal9m-H300R"
P3,"FragMix, Anneal2m-D44V, Anneal6m-Q268R, Anneal7m-E279K, Anneal8, Anneal9"
A4,"FragMix, Anneal2m-D44V, Anneal6m-Q268R, Anneal7m-E279K, Anneal8m-K285Q, Anneal9m-H300R"
B4,"FragMix, Anneal2m-D44V, Anneal6m-Q268R, Anneal7m-E279K, Anneal8, Anneal9m-H300R"
C4,"FragMix, Anneal2, Anneal6, Anneal7m-E279K, Anneal8, Anneal9"
D4,"FragMix, Anneal2, Anneal6, Anneal7, Anneal8, Anneal9"
E4,"FragMix, Anneal2m-D44V, Anneal6m-Q268R, Anneal7m-E279K, Anneal8, Anneal9"
F4,"FragMix, Anneal2m-D44V, Anneal6m-Q268R, Anneal7m-E279K, Anneal8, Anneal9"
G4,"FragMix, Anneal2, Anneal6, Anneal7, Anneal8, Anneal9"
H4,"FragMix, Anneal2, Anneal6m-Q268R, Anneal7m-E279K, Anneal8m-K285Q, Anneal9m-H300R"
I4,"water1, water2, water3, water4, water5"
J4,"FragMix, Anneal2m-D44V, Anneal6m-Q268R, Anneal7, Anneal8, Anneal9"
K4,"FragMix, Anneal2m-D44V, Anneal6m-Q268R, Anneal7m-E279K, Anneal8, Anneal9m-H300R"
L4,"FragMix, Anneal2, Anneal6, Anneal7m-E279K, Anneal8, Anneal9"
M4,"FragMix, Anneal2, Anneal6, Anneal7, Anneal8, Anneal9"
N4,"FragMix, Anneal2, Anneal6, Anneal7, Anneal8, Anneal9m-H300R"
O4,"FragMix, Anneal2m-D44V, Anneal6m-Q268R, Anneal7m-E279K, Anneal8m-K285Q, Anneal9m-H300R"
P4,"FragMix, Anneal2, Anneal6, Anneal8, Anneal9"
A5,"FragMix, Anneal2m-D44V, Anneal6m-Q268R, Anneal7m-E279K, Anneal8m-K285Q, Anneal9"
B5,"FragMix, Anneal2, Anneal6, Anneal7, Anneal8m-K285Q, Anneal9"
C5,"FragMix, Anneal2m-D44V, Anneal6m-Q268R, Anneal7, Anneal8m-K285Q, Anneal9m-H300R"
D5,"FragMix, Anneal2m-D44V, Anneal6m-Q268R, Anneal7, Anneal8, Anneal9"
E5,"FragMix, Anneal2m-D44V, Anneal6m-Q268R, Anneal7m-E279K, Anneal8, Anneal9"
F5,"FragMix, Anneal2m-D44V, Anneal6, Anneal7m-E279K, Anneal8m-K285Q, Anneal9m-H300R"
G5,"FragMix, Anneal2, Anneal6m-Q268R, Anneal7m-E279K, Anneal8m-K285Q, Anneal9m-H300R"
H5,"FragMix, Anneal2m-D44V, Anneal6, Anneal7m-E279K, Anneal8m-K285Q, Anneal9m-H300R"
I5,"FragMix, Anneal2m-D44V, Anneal6, Anneal7m-E279K, Anneal8m-K285Q, Anneal9m-H300R"
J5,"FragMix, Anneal2m-D44V, Anneal6, Anneal7m-E279K, Anneal8m-K285Q, Anneal9m-H300R"
K5,"FragMix, Anneal2, Anneal6, Anneal7, Anneal8, Anneal9"
L5,"FragMix, Anneal2m-D44V, Anneal6m-Q268R, Anneal7, Anneal8m-K285Q, Anneal9m-H300R"
M5,"water1, water2, water3, water4, water5"
N5,"FragMix, Anneal2, Anneal6m-Q268R, Anneal7, Anneal8, Anneal9"
O5,"FragMix, Anneal2, Anneal6, Anneal7, Anneal8, Anneal9m-H300R"
P5,"FragMix, Anneal2m-D44V, Anneal6m-Q268R, Anneal7m-E279K, Anneal8m-K285Q, Anneal9m-H300R"
A6,"FragMix, Anneal2, Anneal6, Anneal7, Anneal8, Anneal9"
B6,"FragMix, Anneal2, Anneal6, Anneal7, Anneal8, Anneal9m-H300R"
C6,"FragMix, Anneal2m-D44V, Anneal6, Anneal7, Anneal8, Anneal9"
D6,"FragMix, Anneal2m-D44V, Anneal6, Anneal7, Anneal8, Anneal9"
E6,"FragMix, Anneal2, Anneal6, Anneal8, Anneal9"
F6,"FragMix, Anneal2, Anneal6m-Q268R, Anneal7m-E279K, Anneal8m-K285Q, Anneal9m-H300R"
G6,"FragMix, Anneal2, Anneal6m-Q268R, Anneal7, Anneal8, Anneal9"
H6,"FragMix, Anneal2, Anneal6, Anneal7, Anneal8, Anneal9"
I6,"water1, water2, water3, water4, water5"
J6,"FragMix, Anneal2m-D44V, Anneal6m-Q268R, Anneal7m-E279K, Anneal8, Anneal9m-H300R"
K6,"FragMix, Anneal2m-D44V, Anneal6, Anneal7, Anneal8, Anneal9"
L6,"FragMix, Anneal2m-D44V, Anneal6m-Q268R, Anneal7m-E279K, Anneal8, Anneal9m-H300R"
M6,"FragMix, Anneal2m-D44V, Anneal6m-Q268R, Anneal7m-E279K, Anneal8m-K285Q, Anneal9m-H300R"
N6,"FragMix, Anneal2m-D44V, Anneal6m-Q268R, Anneal7m-E279K, Anneal8m-K285Q, Anneal9"
O6,"FragMix, Anneal2, Anneal6, Anneal7m-E279K, Anneal8, Anneal9"
P6,"FragMix, Anneal2, Anneal6m-Q268R, Anneal7, Anneal8, Anneal9"
'''

fragFileCSV = '''Frag ID,Plate,Well
Anneal2,storage_plate1,A1
Anneal6,storage_plate1,B1
Anneal7,storage_plate1,C1
Anneal8,storage_plate1,D1
Anneal9,storage_plate1,E1
Anneal2m-D44V,storage_plate1,F1
Anneal6m-Q268R,storage_plate1,G1
Anneal7m-E279K,storage_plate1,H1
Anneal8m-K285Q,storage_plate1,I1
Anneal9m-H300R,storage_plate1,J1
FragMix,storage_plate1,K1
water1,storage_plate1,L1
water2,storage_plate1,M1
water3,storage_plate1,N1
water4,storage_plate1,O1
water5,storage_plate1,P1
'''

# protocol run function
def run(protocol: protocol_api.ProtocolContext):
    def droptip(pipette, testing = False):
        if testing == False:
            pipette.drop_tip()
        else:
            pipette.blow_out(protocol.fixed_trash["A1"])
            pipette.return_tip()
    # Specify if you are testing with food coloring or not:
    testing = False
    trash_testing = not testing
    Temperature = not testing # True or False only
    # Change adjustment to 384 well plate if you are only using part of it. 
    # For example, set to 12 if using right half of the plate and 0 if left.
    adjustment_384 = 0
    
    assembly_plate = protocol.load_labware(
        "appliedbiosystemsmicroamp_384_wellplate_40ul", location="6"
    ) # Assembled DNA fragments from Majeste's Golden Gate Assembly
    reaction_plate = protocol.load_labware(
        "appliedbiosystemsmicroamp_384_wellplate_40ul", location="3"
    ) # Final plate where reaction is assembled
    tiprack = protocol.load_labware(
        "opentrons_96_tiprack_20ul", location="2"
    ) # 20uL tips for the premade solution transfer from premade to PCR
    tiprack2 = protocol.load_labware(
        "opentrons_96_tiprack_20ul", location="1"
    ) # 20uL tips for the DNA transfer from GGA "DNA" to PCR
    tiprack3 = protocol.load_labware(
        "opentrons_96_tiprack_20ul", location="7"
    ) # 20uL tips for the DNA transfer from GGA "DNA" to PCR
    tiprack4 = protocol.load_labware(
        "opentrons_96_tiprack_20ul", location="8"
    ) # 20uL tips for the DNA transfer from GGA "DNA" to PCR
    tiprack5 = protocol.load_labware(
        "opentrons_96_tiprack_20ul", location="9"
    ) # 20uL tips for the DNA transfer from GGA "DNA" to PCR
    # Load reagent plate on temperature module to keep it cool
    temp_mod = protocol.load_module(
        module_name="temperature module gen2", location="4"
    )
    reagent_plate = temp_mod.load_labware(
        "opentrons_96_aluminumblock_generic_pcr_strip_200uL"
    )
    # Pre-made solution of Q5 MasterMix, Forward & Reverse Primers, and Nuclease-Free Water
    
    # pipettes
    left_pipette = protocol.load_instrument(
        "p20_multi_gen2", mount="left",
        tip_racks=[tiprack,tiprack2,tiprack3,tiprack4,tiprack5]
    )
    
    right_pipette = protocol.load_instrument(
        "p20_single_gen2", mount="right",
        tip_racks=[tiprack,tiprack2,tiprack3,tiprack4,tiprack5]
    )
    
    if testing == False:
        temp_mod.set_temperature(celsius=12) # sets a standard temperature while the other components are being mixed/added
    
    # Wells
    # Download fragment/sequence wells from CSV files
    seqFile = pd.read_csv(io.StringIO(seqFileCSV)) # Imports CSV as pandas dataframe
    fragFile = pd.read_csv(io.StringIO(fragFileCSV))
    seqFileData = seqFile.values.tolist() # Converts pandas dataframe to list of list
    fragFileData = fragFile.values.tolist()
    # Specify plate volumes, dead volumes, and fragment volumes
    fragVolume = 2
    maxWellVolume = 240
    deadVolume = 35
    # Get aspirate speeds
    default_right_aspirate_rate = right_pipette.flow_rate.aspirate # ul/s
    default_right_dispense_rate = right_pipette.flow_rate.dispense # ul/s
    
    # Find active columns and add to list
    active_columns = []
    active_column_numbers = []
    for seqrow in seqFileData:
        wellID = seqrow[0].lstrip(" ")
        wellID = wellID.rstrip(" ")
        column_number = wellID[1:]
        column_ID = "A" + column_number
        if reaction_plate[column_ID] not in active_columns:
            active_columns.append(reaction_plate["A" + column_number])
            active_columns.append(reaction_plate["B" + column_number])
            active_column_numbers.append(column_number)
    
    # Divide columns for distribute by the maximum amount of volume held in reagent well
    reagent_well_volume = 200
    GGA_deadVolume = 10
    GGA_reagent_volume = 2.5
    divided_columns = []
    current_column = []
    reagent_counter = 1
    max_counter = math.floor((reagent_well_volume - GGA_deadVolume) / GGA_reagent_volume)        
    for index in range(len(active_columns)):
        if index == len(active_columns) - 1:
            current_column.append(active_columns[index])
            divided_columns.append(current_column)
        else:
            if reagent_counter <= max_counter:
                current_column.append(active_columns[index])
                reagent_counter += 1
            else:
                divided_columns.append(current_column)
                current_column = [active_columns[index]]
                reagent_counter = 2
            
    # Generate source/reagent wells corresponding to divided columns
    reagent_source_wells = reagent_plate.rows()[0][:len(divided_columns)]
    
    # Assemble fragment mixtures:
    # This function generates a data structure I can use for fragment assembly in OT-2
    def GetFragmentWells(seqFileData, fragVolume, maxWellVolume, deadVolume):
        Wells384 = []
        fragIDList = []
        fragWellUsages = []
        fragPlateIDs = []
        fragPlateWells = []
        fragPlateVolumes = []
        fragPlateAssignments = []
        # Generate well IDs for 384 well plate
        for column in range(1,25):
            for row in ["A", "B", "C", "D", "E", "F", "G", "H", "I", "J", "K", "L", "M", "N", "O", "P"]:
                Wells384.append(row + str(column))
        # Get all the fragment IDs and how many wells they are used in
        for seqrow in seqFileData:
            fragList = seqrow[1].split(",")
            # strip any leading/trailing white space
            for index in range(len(fragList)):
                fragList[index] = fragList[index].lstrip(" ")
                fragList[index] = fragList[index].rstrip(" ")
                if fragList[index] not in fragIDList:
                    fragIDList.append(fragList[index])

        # Get number of wells each fragment is used in
        for fragID in fragIDList:
            fragUsage = 0
            for seqrow in seqFileData:
                fragList = seqrow[1].split(",")
                # strip any leading/trailing white space
                for index in range(len(fragList)):
                    fragList[index] = fragList[index].lstrip(" ")
                    fragList[index] = fragList[index].rstrip(" ")
                if fragID in fragList:
                    fragUsage += 1
            fragWellUsages.append([fragID,fragUsage])

        # Sort fragWellUsages by usage:
        fragWellUsages.sort(key = lambda x: x[1])
        fragWellUsages.reverse()

        # Go through all the fragIDs to generate their well assignments
        for fragID in fragWellUsages:
            fragPlateIDs.append(fragID[0])
            fragDestinationWells = []
            fragPlateWells.append(Wells384[0])
            Wells384.pop(0) # Pop the element so it is no longer assigned
            for seqrow in seqFileData:
                fragList = seqrow[1].split(",")
                # strip any leading/trailing white space
                for index in range(len(fragList)):
                    fragList[index] = fragList[index].lstrip(" ")
                    fragList[index] = fragList[index].rstrip(" ")
                if fragID[0] in fragList:
                    fragDestinationWells.append(seqrow[0])
                if len(fragDestinationWells) > math.floor((maxWellVolume-deadVolume-fragVolume)/fragVolume) :
                    fragPlateVolumes.append(len(fragDestinationWells)*fragVolume + deadVolume)
                    fragPlateIDs.append(fragID[0])
                    fragPlateVolumes.append(len(fragDestinationWells)*fragVolume + deadVolume)
                    fragPlateAssignments.append([fragPlateWells[-1]]+fragDestinationWells)
                    fragPlateWells.append(Wells384[0])
                    Wells384.pop(0) # Pop the element so it is no longer assigned
                    fragDestinationWells = []

            fragPlateVolumes.append(len(fragDestinationWells)*fragVolume + deadVolume)
            fragPlateAssignments.append([fragPlateWells[-1]]+fragDestinationWells)
        
        return fragPlateAssignments
    
    def dispenseHeight(well):
        wellDepth = 9.09 # mm
        wellVolume  = 40 # μL
        adjustmentHeight = math.ceil((fragVolume/wellVolume) * wellDepth * 10) / 10
        height = 0.75 + fragWellDispenses[well] * adjustmentHeight
        return height

    fragAssignments = GetFragmentWells(seqFileData, fragVolume, maxWellVolume, deadVolume)
    fragWellDispenses = {} # Tracks the number of fragments added to well to adjust dispense height
    # Initialize every well in 384 plate
    for column in range(1,25):
        for letter in list(string.ascii_uppercase[0:16]):
            fragWellDispenses[letter+str(column)] = 0

    # Count dispenses during fragment mixing
    for fragAssignment in fragAssignments:
        # Sort wells to dispense in the ones with the smallest number of previous fragments first to minimize contamination
        fragWellDispenses = dict(sorted(fragWellDispenses.items(), key=lambda item: item[1]))
        destination_wells = []
        for key in fragWellDispenses.keys():
            if key in fragAssignment[1:]:
                destination_wells.append(assembly_plate[key].bottom(dispenseHeight(key)))
                fragWellDispenses[key] += 1

    # Distribute GGA reagent/master mix into wells of reaction plate
    golden_gate_pipette_height = 0.5 # mm
    left_pipette.pick_up_tip()
    for index in range(len(divided_columns)):
        left_pipette.mix(4,20,reagent_source_wells[index].bottom(golden_gate_pipette_height)) # Mix before
        dest_wells = []
        for well in divided_columns[index]:
            dest_wells.append(well.bottom(golden_gate_pipette_height))
        left_pipette.distribute(
            volume = GGA_reagent_volume,
            source = reagent_source_wells[index].bottom(golden_gate_pipette_height),
            dest = dest_wells,
            new_tip = "never",  # "once", "always", or None will error
            trash = trash_testing,
            disposal_volume = 0
        )
    
    droptip(left_pipette,testing)
    
    # Add DNA to the wells in duplicate
    fragment_pipette_height = 0.5 # mm
    fragMixVolume = fragWellDispenses[seqFileData[0][0]]*fragVolume/2
    for column in active_column_numbers:
        left_pipette.pick_up_tip()
        left_pipette.mix(3,fragMixVolume,assembly_plate["A" + column].bottom(fragment_pipette_height)) # Mix PCR products
        left_pipette.aspirate(3.5, assembly_plate["A" + column].bottom(fragment_pipette_height))
        left_pipette.dispense(3.5, reaction_plate["A" + column].bottom(fragment_pipette_height))
        left_pipette.mix(6,5,reaction_plate["A" + column].bottom(fragment_pipette_height)) # Mix reactions
        droptip(left_pipette,testing)
    for column in active_column_numbers:
        left_pipette.pick_up_tip()
        left_pipette.mix(3,fragMixVolume,assembly_plate["B" + column].bottom(fragment_pipette_height)) # Mix PCR products
        left_pipette.aspirate(3.5, assembly_plate["B" + column].bottom(fragment_pipette_height))
        left_pipette.dispense(3.5, reaction_plate["B" + column].bottom(fragment_pipette_height))
        left_pipette.mix(6,5,reaction_plate["B" + column].bottom(fragment_pipette_height)) # Mix reactions
        droptip(left_pipette,testing)
```
