## Supporting FIle 1 for "Rapid Cell-Free Combinatorial Mutagenesis Workflow Using Small Oligos Suitable for High-Iteration, Active Learning-Guided Protein Engineering": UricaseAssayScript.py

```
from opentrons import protocol_api

metadata = {
    'protocolName': 'Uricase Normalization Assay',
    'author': 'R. Godin w/ChatGPT',
    'description': """Distribute cell-free product into 384-well plates and set up uric acid assay 
    and splitGFP normalization assay with them""",
    'apiLevel': '2.15'
}

# Global testing flag
testing = False  # Set to False to drop tips into trash

# Conditional tip drop function
def droptip(pipette):
    if not testing:
        pipette.drop_tip()
    else:
        pipette.blow_out(pipette.trash_container.wells()[0])
        pipette.return_tip()

def run(protocol: protocol_api.ProtocolContext):

    # Labware
    cell_free_plate = protocol.load_labware('corning_384_wellplate_50ul', '1', label='Cell-Free Plate')
    uricase_plate = protocol.load_labware('corning_384_wellplate_112ul_flat', '2', label='Uricase Plate')
    splitGFP_plate = protocol.load_labware('corning_384_wellplate_112ul_flat', '3', label='splitGFP Plate')
    shared_reservoir = protocol.load_labware('nest_12_reservoir_15ml', '4', label='Shared Reservoir')

    # Tip racks
    tiprack_p20 = protocol.load_labware('opentrons_96_filtertiprack_20ul', '5')
    tiprack_p300 = protocol.load_labware('opentrons_96_tiprack_300ul', '6')

    # Pipettes
    p20 = protocol.load_instrument('p20_multi_gen2', mount='left', tip_racks=[tiprack_p20])
    p300 = protocol.load_instrument('p300_multi_gen2', mount='right', tip_racks=[tiprack_p300])

    # Reagents from shared reservoir
    uric_acid = shared_reservoir.wells_by_name()['A1']
    splitGFP_reagent = shared_reservoir.wells_by_name()['A12']

    # Parameters
    start_col = 1
    end_col = 6
    aspiration_height = 0.5  # mm from bottom
    p300_aspiration_height = -14 # mm from bottom
    dispense_height = 0.2    # mm from bottom for cell-free stamping

    # === STEP 1 === Transfer 6 µL from cell_free_plate to uricase & splitGFP
    original_aspirate_rate = p20.flow_rate.aspirate
    original_dispense_rate = p20.flow_rate.dispense

    p20.flow_rate.aspirate = original_aspirate_rate * 0.25
    p20.flow_rate.dispense = original_dispense_rate * 0.25

    for col in range(start_col, end_col + 1):
        col_idx = col - 1
        for row_offset in [0, 1]:  # rows A and B
            source_well = cell_free_plate.columns()[col_idx][row_offset]
            splitGFP_well = splitGFP_plate.columns()[col_idx][row_offset]
            uricase_well = uricase_plate.columns()[col_idx][row_offset]

            p20.pick_up_tip()
            p20.mix(5, 5, source_well.bottom(aspiration_height))
            p20.aspirate(7, source_well.bottom(aspiration_height))

            p20.dispense(4, uricase_well.bottom(dispense_height))
            p20.touch_tip(uricase_well, v_offset=-11.3, speed=20)

            p20.dispense(2, splitGFP_well.bottom(dispense_height))
            p20.touch_tip(splitGFP_well, v_offset=-11.3, speed=20)

            droptip(p20)

    # Reset to default rates
    p20.flow_rate.aspirate = original_aspirate_rate
    p20.flow_rate.dispense = original_dispense_rate

    # === STEP 2 === Uric acid: aspirate 300 µL, dispense 96 µL × 2, blow out at source.top()
    p300.pick_up_tip()
    for col in range(start_col, end_col + 1):
        col_idx = col - 1
        dest_wells = [uricase_plate.columns()[col_idx][row_offset] for row_offset in [0, 1]]
        p300.aspirate(200, uric_acid.bottom(p300_aspiration_height))
        for dest in dest_wells:
            p300.dispense(96, dest.top(-5))
        p300.blow_out(uric_acid.top())
    droptip(p300)

    # === STEP 3 === splitGFP: aspirate 240 µL, dispense 78 µL × 2, blow out at source.top()
    p300.pick_up_tip()
    for col in range(start_col, end_col + 1):
        col_idx = col - 1
        dest_wells = [splitGFP_plate.columns()[col_idx][row_offset] for row_offset in [0, 1]]
        p300.aspirate(160, splitGFP_reagent.bottom(p300_aspiration_height))
        for dest in dest_wells:
            p300.dispense(78, dest.top(-5))
        p300.blow_out(splitGFP_reagent.top())
    droptip(p300)
```
